# Sensing volatiles throughout the body: Geographic and tissue-specific olfactory receptor expression in the fig wasp *Ceratosolen fusciceps*

**DOI:** 10.1101/2023.03.03.530950

**Authors:** Sushma Krishnan, Snehal Dilip Karpe, Hithesh Kumar, Lucy B Nongbri, Sowdhamini Ramanathan, Ewald Grosse-Wilde, Bill S. Hansson, Renee M. Borges

## Abstract

An essential adaptive strategy in insects is the evolution of olfactory receptors (ORs) to recognize important volatile environmental chemical cues. Our model species, *Ceratosolen fusciceps,* a specialist wasp pollinator of *Ficus racemosa*, likely possesses an OR repertoire that allows it to distinguish fig-specific volatiles in highly variable environments. Using a newly assembled genome-guided transcriptome, we annotated 63 ORs in the species and reconstructed the phylogeny of *Ceratosolen* ORs in conjunction with other hymenopteran species. Expression analysis showed that though ORs were mainly expressed in the antennae, 20 percent were also expressed in non-antennal tissues such as the head, thorax, abdomen, legs, wings, and ovipositor. Specific upregulated expression was observed in OR30C in the head and OR60C in the wings. We identified OR expression from all major body parts of *C. fusciceps*, suggesting novel roles of ORs throughout the body. Further examination of OR expression of *C. fusciceps* in widely separated geographical locations, i.e., south (urban) and northeast (rural) India, revealed distinct OR expression levels in different locations. This discrepancy likely parallels the observed variation in fig volatiles between these regions and provides new insights into the evolution of insect ORs and their expression across geographical locations and tissues.

## Introduction

Chemical interactions between plant volatiles and insect olfactory receptors (ORs) are essential processes for the survival and reproduction of insects. In insects, ORs are expressed in the dendritic membrane of olfactory sensory neurons (OSNs). Insect ORs are 7-transmembrane domain proteins with an intracellular N-terminus and an extracellular C-terminus (Benton et al. 2006; Smart et al. 2008; Missbach et al. 2014). Functional insect ORs consist of heterodimeric complexes with a highly divergent ligand-specific OR and a highly conserved co-receptor (Orco) (Larsson et al. 2004; Sato et al. 2008; Wicher et al. 2008; Butterwick et al. 2018). During the evolution of insects, ORs became a massively diverse gene family as a result of adaptation to complex and changeable chemical environments (McBride 2007; Linz et al. 2013; Schmidt and Benton 2020; Wicher and Miazzi 2021). In the case of specialist insect pollinators, it is reasonable to expect adaptation to the detection of signature host plant chemical signals.

The fig and fig wasp interaction has emerged as an important model in chemical ecology because of its highly specific mutualism (e.g., Bain et al., 2016; Borges et al., 2008; Borges, 2015, 2021; Borges et al., 2011, 2013; Ghara et al., 2011; Hou et al., 2020; Proffit et al., 2007; Ranganathan & Borges, 2009; Wei et al., 2021; Xin et al., 2020; Yadav & Borges, 2017). This interplay also presents these wasps as an attractive model for the study of olfactory adaptations. Agaonid female wasps enter the enclosed globular fig inflorescence called the syconium, oviposit concurrently with pollination, and later die within the syconium. Female wasps get only one chance to enter an oviposition/pollination chamber since they lose their wings and part of their antennae in the process of entering the syconium. Therefore, female fig wasps are selected for high specificity towards the pollination scent composed of volatile organic compounds (VOCs) emitted by the pollen-receptive host fig. Male wasps eclose in the enclosed syconium, where they die after mating with eclosed females and thus do not have to find a mate over a long distance. Probably due to this highly specialized lifestyle, pheromones and their receptors are unknown in fig wasps. Finding a pollen-receptive fig syconium is the key for the female pollinator for oviposition and reproduction. We used the widely distributed *Ficus racemosa* and its wasp pollinator *Ceratosolen fusciceps* for the molecular characterization of ORs in northeastern and southern India since this particular fig and wasp species pair has received considerable attention regarding chemical signaling (Bain et al. 2016; Xiao et al. 2021). When insects are highly specialized on their host, the rate of OR gene function loss is 9-to-10 times greater than in generalists as seen in highly specialized compared to generalized drosophilid flies (McBride 2007). The extreme reduction of OR gene numbers in *Ceratosolen solmsi*, an obligate and host-specific pollinator of *Ficus hispida* (just 56 compared to about ∼100-300 in other solitary parasitic or predatory Apoid wasps) is likely due to its obligate plant host specificity (Obiero et al. 2021), and we expected a similar reduction in *C. fusciceps*.

It was initially hypothesized that ORs were exclusively expressed in olfactory tissues such as antennae and mouth parts. However, *Culex* mosquitoes (Leal et al. 2013), leaf beetles (Wang et al. 2016), green plant bugs (An et al. 2016), and butterflies (van Schooten et al. 2020) showed expression of some ORs in a variety of tissues other than the antenna, viz. legs, wings, head, thorax, and abdomen. In the tobacco budworm, *Heliothis virescens*, lower levels of pheromone-detecting receptors were expressed in tissues such as the proboscis, abdomen, leg, wing, and thorax (Krieger et al. 2004). Widmayer et al. (2009) showed expression of pheromone receptors in ovipositor sensilla of female *H. virescens.* The ovipositor of the noctuid moth (Koutroumpa et al. 2021), grass moth (Xia et al. 2015), and tobacco hornworm (Klinner et al. 2016) also expresses ORs. In *Spodoptera littoralis* chemosensory receptors were identified from mouthparts, legs, and ovipositors (Koutroumpa et al. 2021). The olfactory repertoire of another extreme specialist marine insect, *Clunio marinus*, revealed the possibility of OR expression in legs, genitalia and larval body (Missbach et al. 2020). The expression of ORs outside of antenna and palps is always paralleled by the presence of sensilla (Koutroumpa et al. 2021). Furthermore, Yadav and Borges (2017) showed experimentally that sensilla on fig wasp ovipositors fire in response to fig volatiles, and the ovipositor deflects in response to carbon dioxide puffs, showing clearly that the ovipositor possesses chemosensory abilities. Broad expression of ORs in tissues other than antennae, maxillary palps, and ovipositors hints towards general functional significance in chemical sensing over the insect body. However, there are very few studies on potential functional significance of olfactory detection in other tissues, like the proboscis of *Manduca sexta* in flower humidity perception and nectar foraging (Goyret and Raguso 2006; Havercamp et al. 2016).

Furthermore, intraspecific variation in signal recognition may also be influenced by the complexity of changing environments (Renou and Anton 2020), and floral scents can vary geographically (Skogen et al. 2022). However, variation in OR expression across geographical locations is rarely studied. To understand OR gene expression variation, most comparative transcriptome analyses in insects have been performed between closely related genera or species (Elgar et al. 2018; Guo et al. 2021) or between genders (Athrey et al. 2021; Xu et al. 2021). Examination of intra-specific OR variation is especially important given changing scenarios of signal content, local volatile environments, and ambient conditions.

Here we compare the host-specific variation of OR expression of pollinating fig wasp. We performed a comparative OR gene expression analysis of *C. fusciceps* tissues across two distinct geographical sites (south India and northeast India bordering China) separated by 3000 km (Fig 1). We expected OR variation between these sites as previous work (Kobmoo et al. 2010; Bain et al. 2016) showed that *C. fusciceps* and its host plant *F. racemosa* form a genetically homogeneous population across south-east Asia (including southern China and Thailand), while populations in south India form separate genetic clusters. We identified 63 ORs in which most ORs were expressed in the antennae, while 20% were also expressed in other olfactory tissues. Upon comparison of ORs from the south and northeast Indian wasps, we observed a few ORs that were exclusively expressed in the antennae of wasps in one region. Other ORs showed significant variation in expression levels between regions which might correspond to variation in fig volatile profiles between these regions (Nongbri and Borges, unpub. data). This inter-population variability in OR expression was also found among ORs expressed outside the antennae.

**Figure 1.**
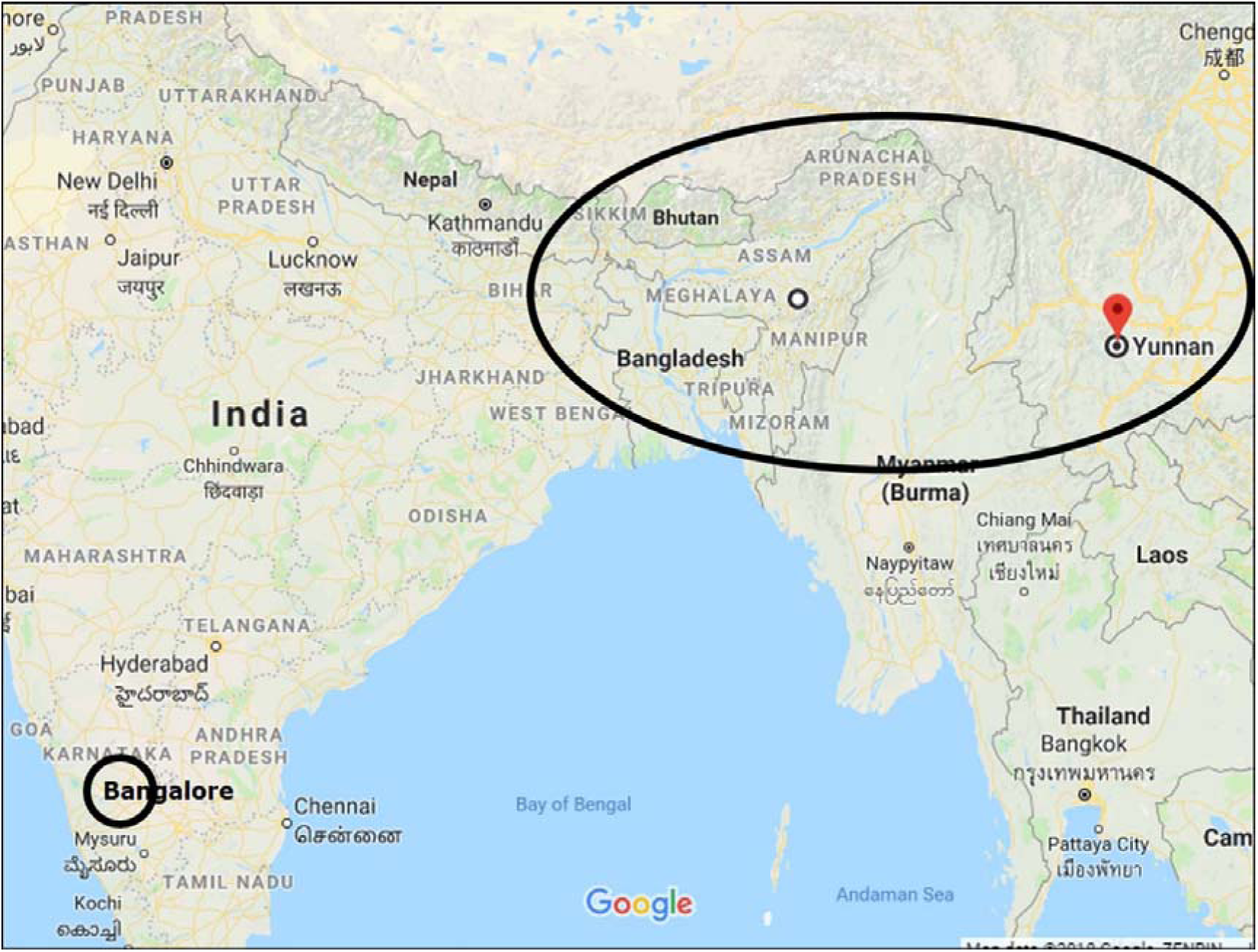
Locations of the study sites in northeastern and southern India. Note that the northeastern site is approximately at the same latitude as the site in Yunnan, China, where earlier data on *F. racemosa* VOCs, plant, and pollinator genetics was established.

## Results

### Total RNA Sequencing and Genome-Guided Transcriptome Assembly

We performed transcriptome sequencing of 14 samples of 7 tissues (antennae, head, abdomen, thorax, legs, wings, and ovipositor) of the fig pollinator wasps collected from 2 different regions (south and northeast India). An average of 23.9 (south India) and 21.2 million (northeast India) preprocessed reads corresponding to an average of 88% of high-quality data of each tissue sample was retained for the assembly (Additional file: Fig. S1). To get the best quality assembly, we assembled the transcriptome sequences by a genome-guided method. For this purpose, we used our newly sequenced and assembled draft genome of *C. fusciceps* (Additional file 1: Table S1). The whole genome of the pollinator wasp was sequenced to get total coverage of 160X using Illumina (100X), Mate pair (40X), and Nanopore (20X) sequencing. We then performed hybrid assembly using MASuRCA (Maryland Super-Read Celera Assembler). As a result, a high-quality genome with a scaffold length of 238 Mb, N50 values of contig 2.2 Mb, and a Scaffold of 4.1 Mb were obtained (Krishnan and Borges, unpublished data). Finally, a good quality transcriptome assembly using Trinity v 2.8.5. was achieved with 58076 transcripts that contained 17746 predicted proteins and 14417 transcripts with known protein domains. BUSCO analysis with Hymenoptera (4415 BUSCOs) and Insecta (1658 BUSCOs) datasets identified 78.2% (3452) and 86.5% (1435) of BUSCOs in our transcriptome assembly (Additional file 1: Table. S2). Based on these results the assembly was considered for further gene annotation.

### Annotation of olfactory receptors in *C. fusciceps*

From the genome-guided transcriptome assembly, we found 74 putative OR transcripts (Additional file 2) using hmmscan against the Pfam database. These results were validated using the InsectOR web server (Karpe et al. 2021); further manual curation resulted in 63 ORs. Among these 63 ORs, 48 sequences encoded for proteins of more than 300 amino acids and were annotated as ‘complete’. Of the remaining sequences 15 transcripts encoded for OR proteins, 8 with missing N-terminus, 3 with missing C-terminus, and 4 captured only the middle fragment of the OR protein sequence. Of the 63 OR protein sequences discovered from the curated transcriptome, 56 had more than 200 amino acid sequence lengths. These were further used for phylogenetic reconstruction of the ORs from two *Ceratosolen* species (*C. solmsi and C. fusciceps*) along with a few other hymenopteran species (Methods). OR sequences containing more than 200 amino acids are generally preferred for phylogenetic analysis (An et al. 2016; Wu et al. 2019; Al-Jalely and Xu 2021; Xu et al. 2021). The inclusion of a diversity of ORs from Hymenoptera also ensured the correct rooting of the tree and its clades. Most of the known hymenopteran OR subfamilies/clades were well-supported in the current reconstruction (Fig 2A, Additional file 1: Table. S3). The ORs of both *Ceratosolen* species had smaller OR repertoires compared to the other hymenopteran species including the two specialized parasitoid wasps, *Microplitis demolitor* (Zhou et al. 2015) and *Nasonia vitripennis* (Robertson et al. 2010). The two *Ceratosolen* species also had the most similar repertoire distribution across OR subfamilies compared to the others.

**Figure 2.**
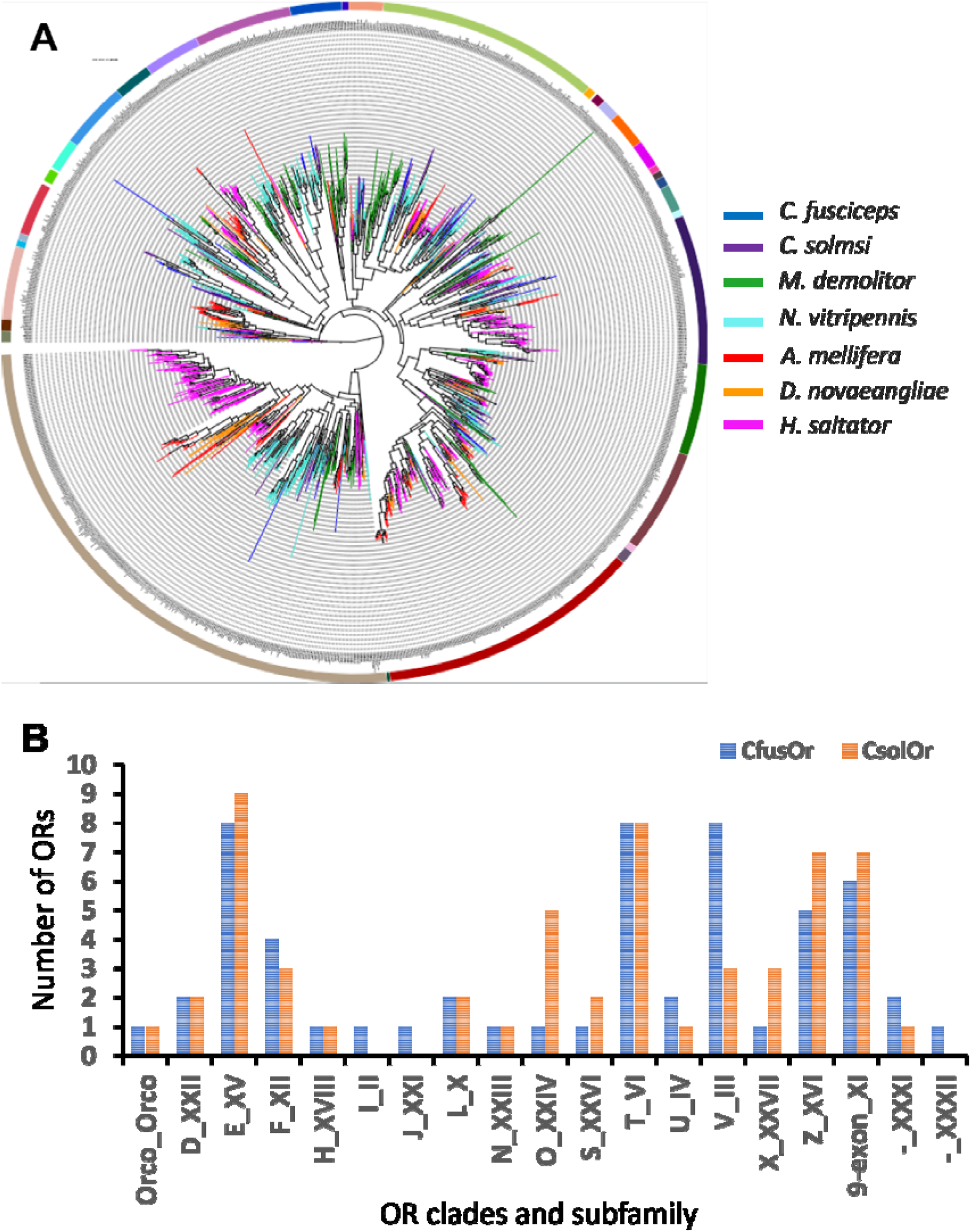
A. Phylogenetic analysis of hymenopteran ORs. The branches were color-coded as per species. The subfamilies are shown by colored stripes. B. Distribution of *C. fusciceps* (CfusOr) and *C. solmsi* (CsolOr) ORs in different clades and subfamilies.

Though *C. fusciceps* ORs were well distributed among hymenopteran clades, striking features were observed for a few OR clades (Fig. 2B, Additional file 1: Table. S3).

The highest number of ORs, 8 each, were observed in clade XV (Subfamily E), clade III (Subfamily V), and clade VI (Subfamily T). The next highest number of ORs 6, 5, and 4 were found in clade XI (subfamily 9-exon), clade XVI (subfamily Z), and clade XII (subfamily F) respectively. Equal distribution was present in clade XXII (subfamily D), clade XXXI (subfamily), clade IV (subfamily U), clade X (subfamily L), and clade X rest (clade X includes Xa and Xb subclades and the remaining ORs that do not form a single clade is named as X rest). The remaining 9 clades had only one OR and 21 clades had no *C. fusciceps* ORs, whereas these clades contained ORs from the other hymenopteran species studied here (Fig. 2B, Additional file 1: Table. S3).

The phylogenetic reconstruction of ORs from the two *Ceratosolen* species (Additional file 1: Fig S2) and the analysis of best-bidirectional BLAST hits demonstrated that although their OR repertoires were similar in size, not all ORs were in a perfect orthologous relationship with *C. solmsi* ORs. Considering both approaches, only 19 OR sequences (ORco, OR-5, 6, 7, 8, 12, 15, 16, 18, 26, 30, 35, 36, 38, 39, 41, 42, 52, 56) had perfect orthologs across the two species and they belong to 6 different OR clades. There were rare species-specific OR expansions, however, and multiple instances of 1:2 or 1:3 protein homologies were found between the two species of *Ceratosolen*.

### OR expression analysis and qPCR validation

A heatmap was generated (Fig 3) from the expression matrix containing transcripts per million (TPM) values (Additional file 3). An OR with TPM > = 1 cutoff was considered for its expression and based on this, of the 63 ORs, 54 were expressed in antennae of south Indian wasps and 9 were not expressed; They are OR28_1, 30C, 51, 54, 55N, 60C, 62, 68, 74F. In northeast Indian wasps, 53 were expressed in antennae and 10 were not expressed. These were OR28_2, 30C, 36, 40F, 54 like_1, 55 like_1F, 60C, 65, 66, 73C. Among the 9 ORs which were not expressed in the antennal tissue from south India, except for OR30C and OR60C, all other 7 ORs were expressed in the antennae of northeast India. Similarly, among the 10 ORs which were not expressed in the antennal tissue from northeast India, except for OR30C and OR60C, all other 8 ORs were expressed in the antennae of south India. Therefore, OR antennal expression in one site did not parallel antennal expression in the other site. In addition, 20% of the antennal ORs from each region showed ectopic tissue expression.

**Figure 3.**
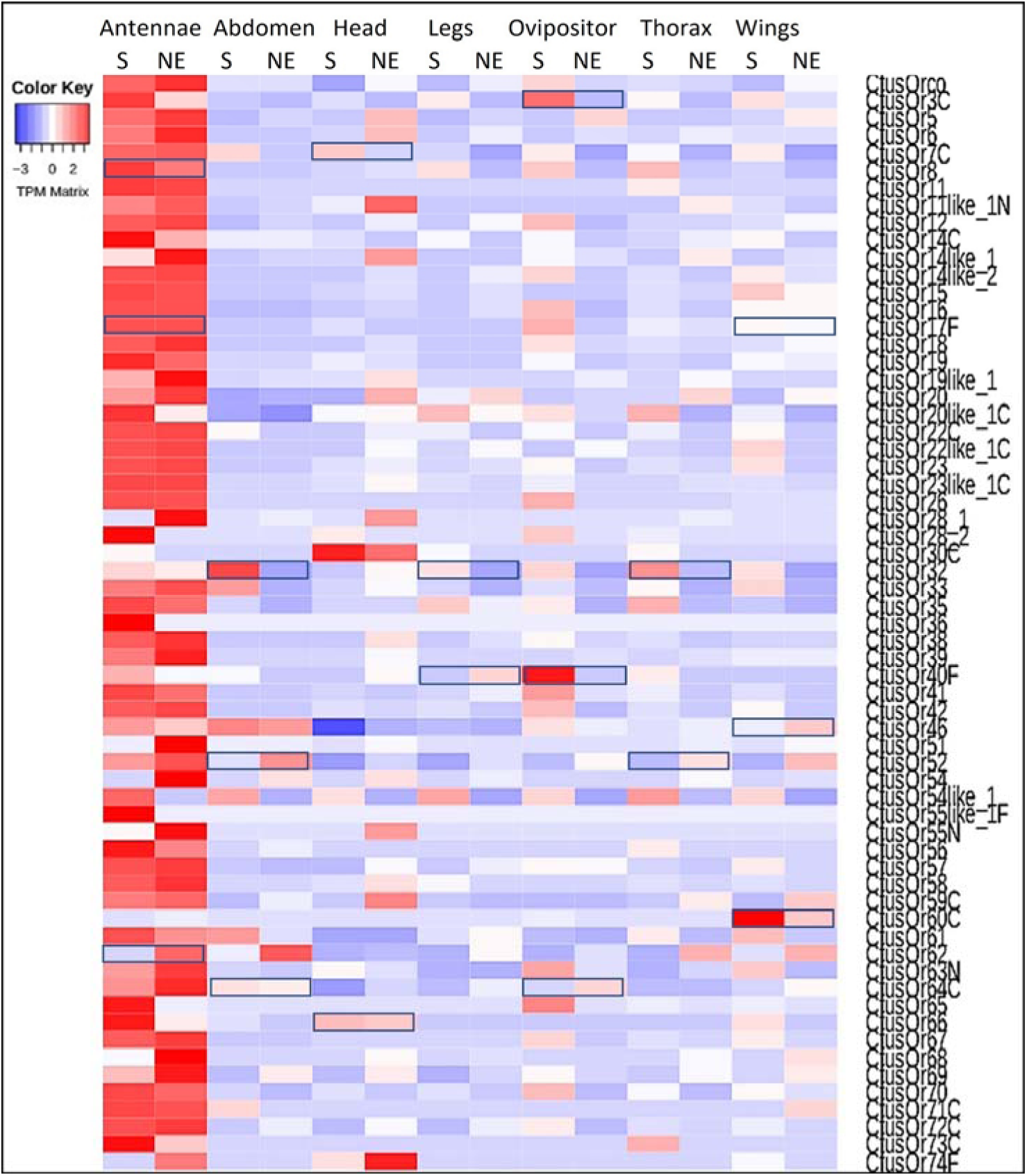
OR expression pattern in *C. fusciceps* tissues across Bangalore (south India) and the northeast Indian region. Heatmap was plotted using heatmap 2 by taking log2 (TPM value +1) and data scaling was done for rows. OR genes indicated in boxes were selected for qPCR analysis and the selection criteria are given in the Methods section.

To confirm the transcription variation observed in the heatmap, we selected a subset of 18 ORs for qPCR analysis (see the justification for selection in Methods, (Fig. 4). They were OR3C, 7C, 8, 12, 17F, 32, 40F, 41, 46, 52, 57, 60C, 62, 64C, 66. These had antennal or non-antennal expressions (Fig. 4).

**Figure 4.**
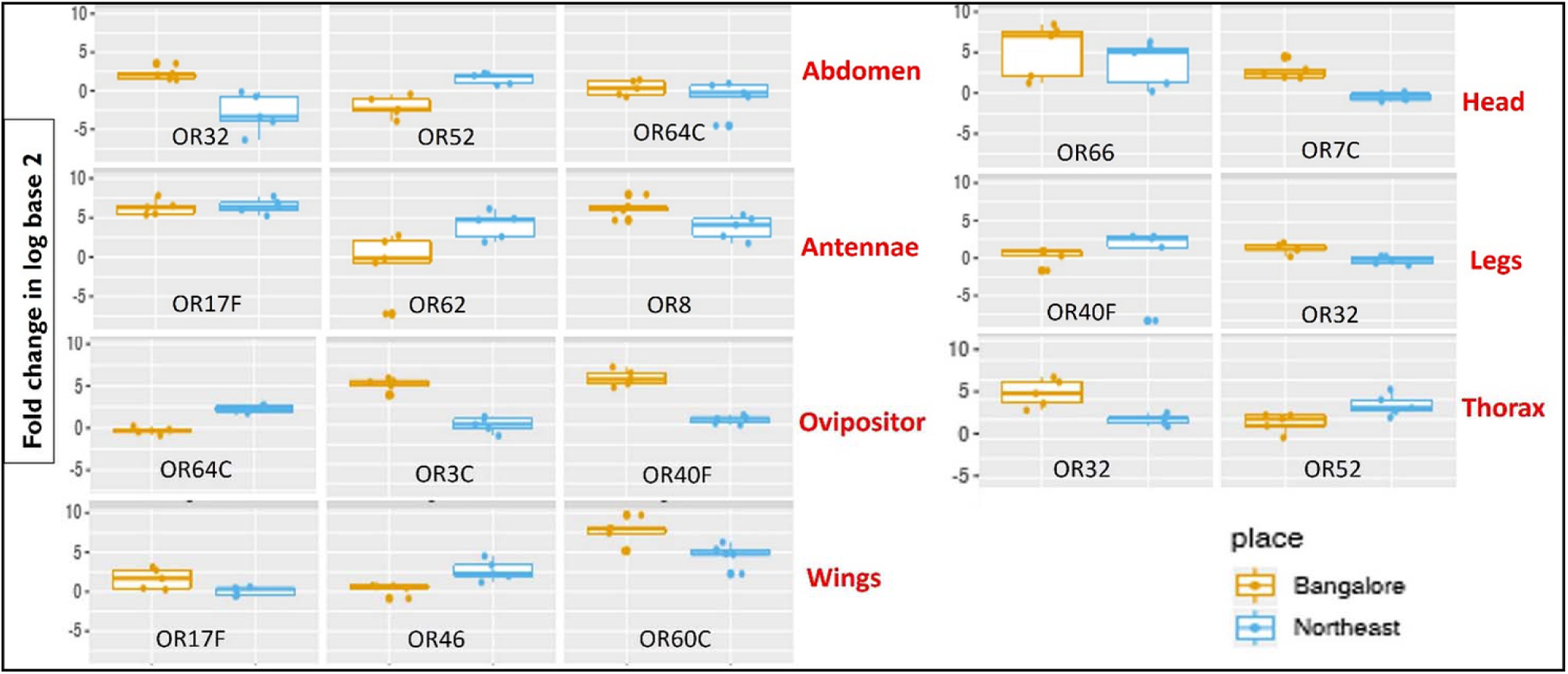
qPCR analysis of *C. fusciceps* ORs from 7 different tissues from south India (Bangalore) and northeast India. Fold-change values in log base 2 values from 5 biological replicates of each tissue were compared across sites by t-tests. P values are provided in Supplemental Table S7. The differential expression of OR genes shown here reproduces the results from the heatmap in Figure 3.

The results of the qPCR analysis showed that OR32 had significantly higher expression in south India than in northeast India, whereas OR52 showed lower expression in south India. OR64C had basal level expression in both regions. In the case of the antennae, OR62 showed a significant increase in expression in northeast samples as compared to south India, whereas OR8 showed the opposite expression pattern. OR17F showed equal expressions in both regions. Similarly, OR7C from the head, OR32 from the legs, OR40F and OR64C from the ovipositor, OR32 and OR52 from the thorax, and OR46, and OR60C from the wings showed significant differences in expression between the two regions. Thus, the qPCR results were in good correspondence with the heatmap derived from TPM values. Therefore, we used the heatmap for further comparison of OR gene expression in different tissues across regions.

### OR expressions were specific to antennal and non-antennal ectopic tissues

For variation across south and northeast India, we depicted the heatmap results as tissue-specific to further analyze OR expression in non-antennal tissues. OR expression is depicted in UpSet plots for south and northeast India (Fig. 5) and these regions combined (Fig. 6).

**Figure 5.**
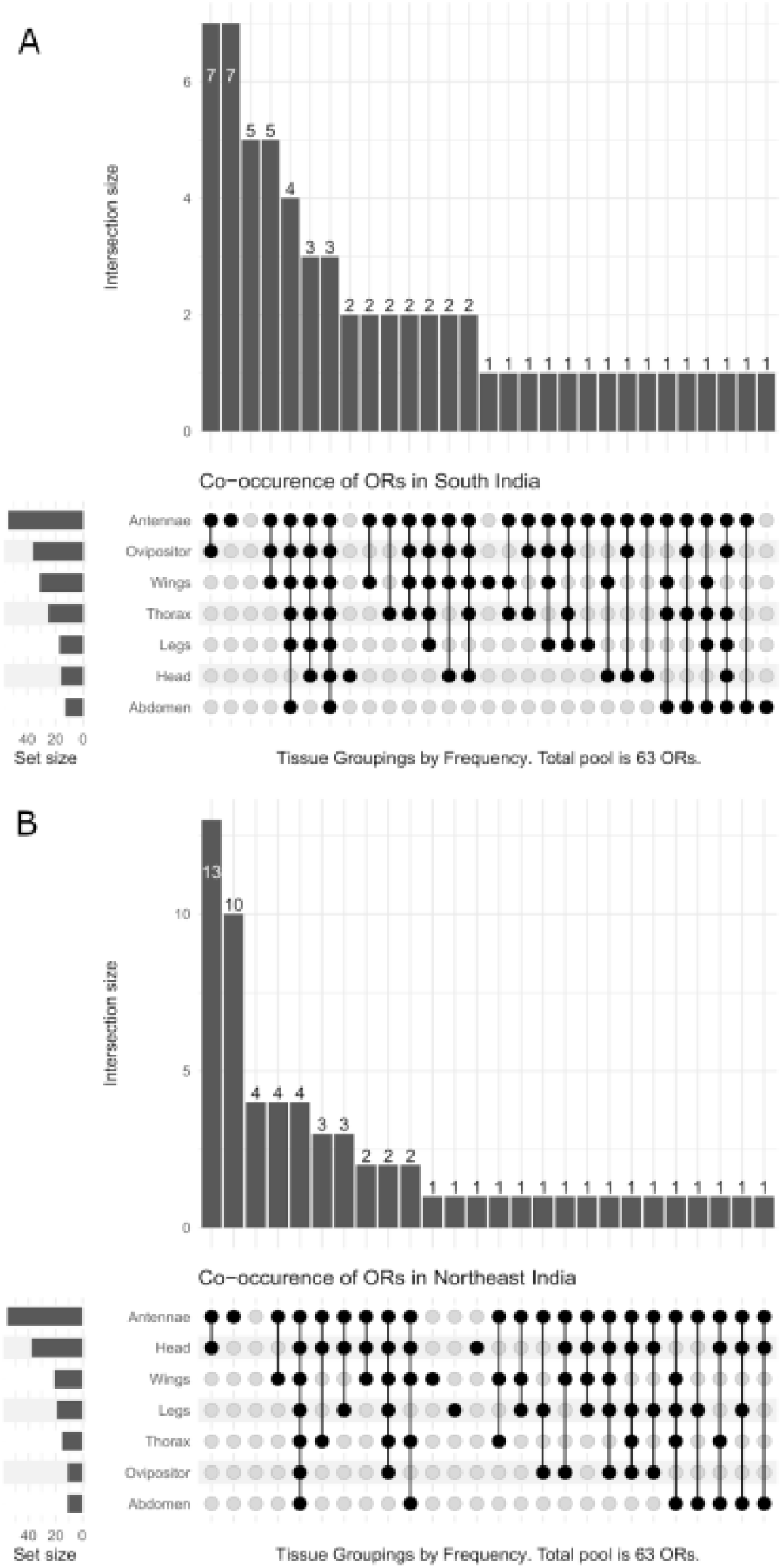
UpSet plots showing tissue-specific OR expression from south and northeast India. A. OR expression profile of south Indian wasps B. OR expression profile of northeast India.

**Figure 6.**
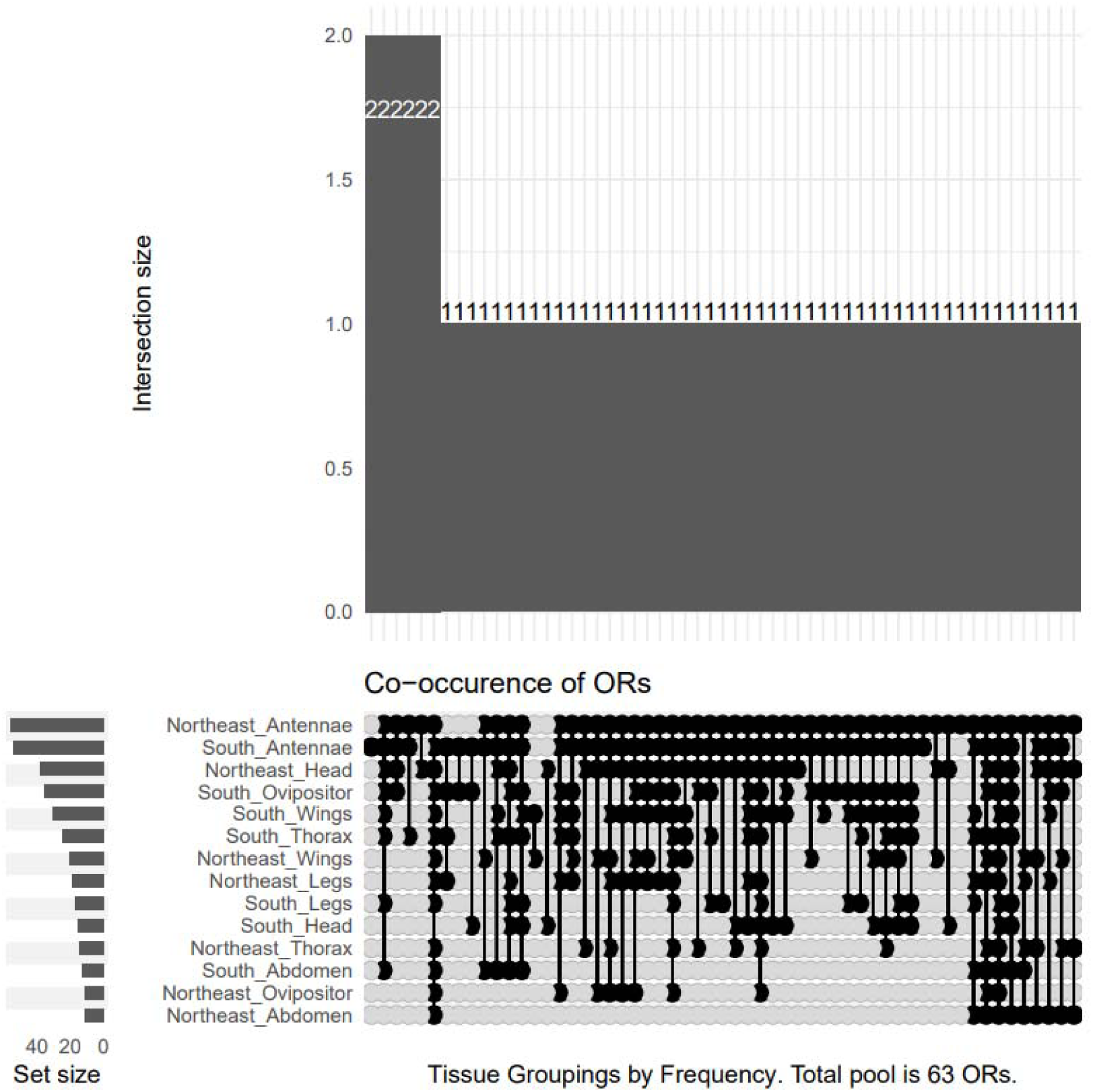
Combined upset plots showing tissue- and region-specific OR expression from south and northeast India.

In the case of south Indian wasps, among 54 antennal OR expressions, 17 ORs were expressed only in the antennae. A total of 10 ORs in the abdomen and legs, 7 ORs in the head, 17 ORs in the thorax, 22 ORs in the wings; 28 ORs in the ovipositor showed expression. Among all these non-antennal ORs, OR30C (XXVII_X subfamily) and OR60C (9 exon subfamily), were expressed specifically in the head and wings respectively. The remaining ORs were expressed in ≥ 2 tissues. OR46, which belongs to the XXIII_N subfamily, and OR54 like_1, which belongs to OR subfamily IV_U, were expressed in all tissues of south Indian wasps (Fig 3, Fig. 5A).

In the case of northeast Indian wasps, among 53 antennal OR expressions, 21 ORs were expressed only in antennae. A total of 8 ORs in the abdomen and thorax, 27 ORs in the head, 11 ORs in the legs, 15 ORs in the wings, and 9 ORs in the ovipositor showed expression. Like the south Indian wasps, OR30C and OR60C were expressed specifically in the head and wings respectively. The remaining ORs were expressed in ≥ 2 tissues. OR46, which belongs to the XXIII_N subfamily, and OR52, which belongs to XV_E, were expressed in all tissues of northeast Indian wasps (Fig 3, Fig. 5B).

### Wasp OR expressions were variable across the region

Among 63 ORs, 46 ORs showed antennal expressions in both south and northeast India; 15 ORs were expressed in only one region; 2 ORs (OR30C and 60C) were not expressed in antennae from any region (Fig 3, Fig 6). Among the 46 antennal OR expressions, 15 ORs showed nearly equal expression levels across regions, whereas the remaining 31 ORs showed moderate to highly variable expressions (Fig 3). This kind of expression variation was also observed in non-antennal tissues such as the abdomen, head, legs, ovipositor, thorax, and wings (Fig 3, Fig. 6). In the abdomen, among the 5 ORs expressed in both regions, 3 ORs showed variable expression. In the head, among 4 ORs expressed in both regions, 2 were variable across regions. In the legs, 5 ORs were expressed in both regions in which 3 ORs were variable. In the ovipositor, 5 ORs were expressed in both the regions in which 4 ORs were variable. In the case of the thorax, 3 ORs were expressed in both regions, and 1 OR was variable in its expression across regions. In the wings, 31 ORs were expressed in both regions and 25 showed variable expressions across regions. OR 46 was expressed in all tissues from both regions.

In both regions, the highest percentage (37% in south India and 43% in northeast India) of ORs were expressed in the antennae (Fig 7). In south India, the next highest level of OR expression was observed from the ovipositor (19%) followed by wings (16%), thorax (12%), legs (6%); the abdomen and head showed the least (5%) OR expression. In northeast India, the next highest level of OR expression was observed from the head (20%) followed by the wings (12%), legs (8%), and abdomen (6%); the thorax, and ovipositor showed the least (5%) OR expression.

**Figure 7.**
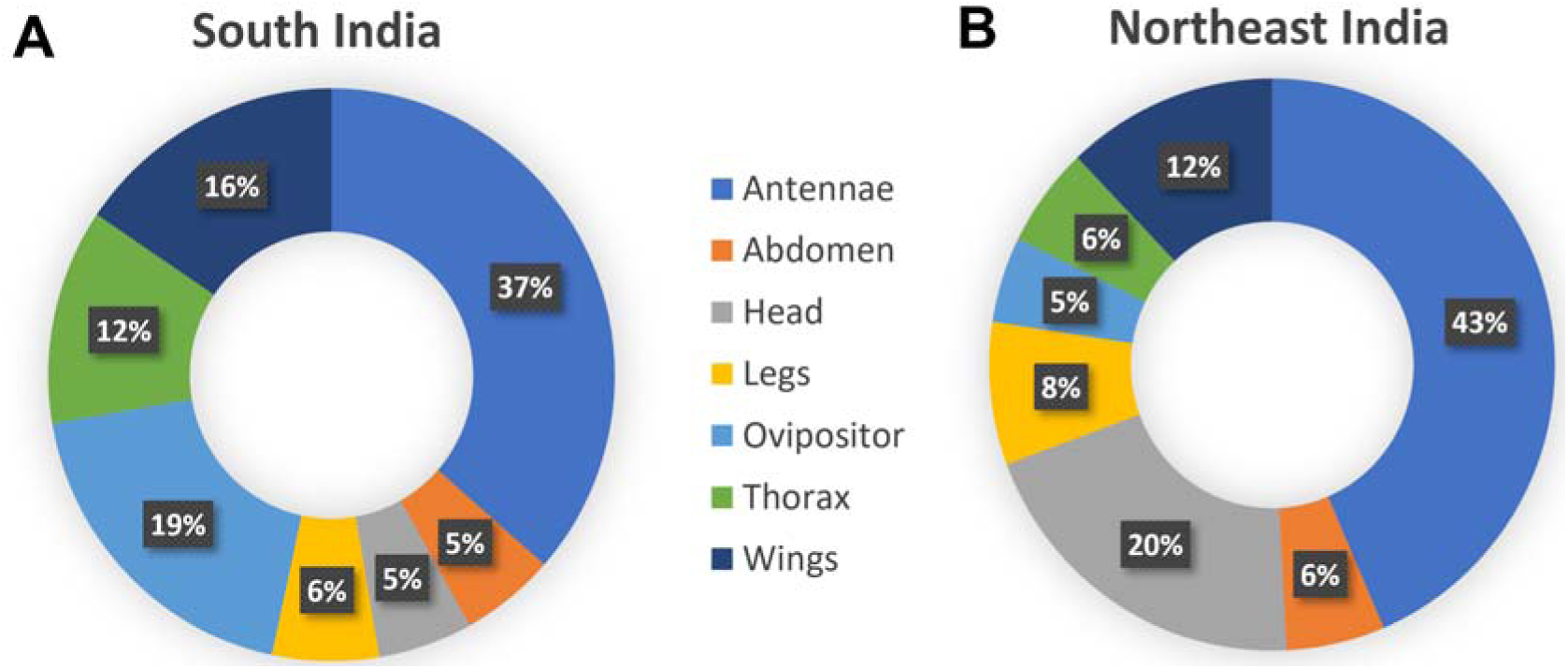
Distribution of OR genes expressed in different fig wasp tissues. A. South India B. Northeast India. The values within the colored sectors are the percentage of ORs expressed in the tissues.

## Discussion

Investigation of OR expression in non-model organisms has unique challenges as well as opportunities. In the present study, we annotated 63 ORs using the newly assembled genome of the non-model organism *C. fusciceps* in India (Krishnan and Borges, unpublished data) and examined differential expression patterns in most tissues of this pollinating wasp. We also compared intraspecific non-chemosensory tissue OR expression patterns across widely separated geographic areas, i.e., south and northeast India. Meanwhile, Xiao et al. (2021) annotated 60 OR transcripts, using the *C. fusciceps* genome of Chinese fig wasp populations that had a scaffold length of 235 Mb, and scaffold N50 of 7.1Mb. The number of genes annotated was also comparable (this study: 12,363; Xiao et al. (2021): 12,171). We were unable to compare the OR sequences from this study with that of the Chinese wasp population because annotated OR sequences for *C. fusciceps* were not provided in Xiao et al. (2021).

The total number of ORs found was comparable to a closely related fig wasp species *C. solmsi* with 56 ORs (Xiao et al. 2013; Zhou et al. 2015). Within the Hymenoptera, the bees and ants investigated for their ORs were mostly generalists; the number of reported ORs is more than 100 for bees (Smith et al. 2011; Karpe et al. 2016, 2017) and more than 200 for ants (Nygaard et al. 2011; Kocher et al. 2015). In wasps, generalist parasitoid wasps such as *N. vitripennis* (Robertson et al. 2010)*, Aenasius bambawale* (Nie et al. 2017), *M. demolitor*, and *Cotesia vestalis* have more than 100 ORs. In specialist fig wasps, less than 100 ORs were recorded (Zhou et al. 2015; Xiao et al. 2021) (Additional file 1: Table S4). This OR reduction may be attributed to the obligate plant host specificity of these wasp species. This hypothesis is further strengthened by the finding that *Drosophila sechellia,* a plant host specialist, is losing ORs 10 times faster than its generalist sibling *D. simulans* (McBride 2007).

Phylogenetic reconstruction shines a light on the differences in OR evolution across the hymenopteran ORs that were compared: i.e., two pollinator fig wasp species *(C. fusciceps and C. solmsi)*, two parasitoid wasps (*N. vitripennis* and *M. demolitor*), a generalist and eusocial honeybee (*Apis mellifera*), a solitary bee that forages specifically (*Dufourea novaeangliae*) and an ant species which is primitively eusocial (*Harpegnathos saltator*) (Fig 2A, Additional file 1: Table S3). Clade XXX (Subfamily Zb) contained expanded ORs from only the two parasitoid wasps and could be important for these two species. Clade XXIII (subfamily N) and clade XXIV (subfamily O) contained one or few ORs per wasp species but none for the other bees and the ant, indicating a probable involvement in a crucial function for the four wasp species but not for the others. Clade IX (subfamily K) was uniquely absent only in the two *Ceratosolen* species but present in low numbers (1 or 2) in the others. Clade XI (subfamily 9-exon) has several cuticular hydrocarbon (CHC)-sensing ORs in the ant species (however, it is not the only CHC sensing OR subfamily) and this was heavily expanded in almost all ant species. Given the CHC-based nest-mate recognition observed in ants (Lorenzi and D’Ettorre 2020), it was predicted that the few ORs from other species belonging to this clade might also recognize CHCs. It was interesting to note that these ORs were expanded to a greater extent in other species compared to the two *Ceratosolen* species. If indeed these receptors are important for CHC detection in all the species, their reduction in *Ceratosolen* could be attributed to the specialization of the two *Ceratosolen* species to specific fig species and the closed chemical environment within which CHC recognition for processes such as mating occurs within the fig syconium. The clades VII (subfamily M) and VIII (subfamily P) were absent in any fig wasp ORs. Subgroup a of clade X (subfamily L) contains the 9-ODA receptor, a receptor for one of the major components of honeybee queen mandibular pheromone (Wanner et al. 2007) and previously identified to be expanded in two eusocial honeybees while almost absent in two solitary bees (Karpe et al. 2017). Therefore, the other ORs from these clades are likely involved in the recognition of the other major components of the honeybee queen mandibular pheromone. As expected, the two *Ceratosolen* species did not display any representative ORs belonging to subgroups Xa and Xb.

Our phylogenetic analysis of ORs from *C. fusciceps* and *C. solmsi* showed that only 19 OR sequences had perfect orthologs across the two species. This was unlike another example of two evolutionarily closely related hymenopteran species of *Apis*, which showed that around 70% of the ORs from the two species had perfect bidirectional best Blast hits as well as phylogenetic clustering (Karpe et al. 2016). Some differences could be associated with the difference in the approach used between this study and Xiao et al. (2013) to annotate the ORs across the two *Ceratosolen* species (from transcriptome vs genome). However, as our transcriptome curation was guided by a high-quality draft genome assembly (Additional file 1: Table S1) and 98% of the transcripts were aligned to the genome, this is an unlikely possibility. The observations of subtle differences in OR evolution across the two species could result from the difference in the volatile profile of their host (*Ficus racemosa* vs *Ficus hispida*) or other environmental forces.

Understanding variation in the insect olfactory system based on variation in the ecologically relevant chemical environment via OR tuning and changes in antennal morphology are already well-established fields. However, possible variation in non-antennal ectopic OR expression and its functional significance is an emerging field of research. OR expression in non-antennal tissues was observed in other insects, e.g. 13 ORs showed expression in non-antennal tissues of the hemipteran *Apolygus lucorum* with 8 ORs exhibiting high expression in heads, legs, and wings (An et al. 2016). OR22 was highly expressed in non-antennal tissues in the coleopteran *Holotrichia oblita* (Li et al. 2017). Expression of the OR2 gene in ectopic tissues of female fig wasps associated with *F. hispida* including the pollinator, *C. solmsi*, strongly indicates the presence of cryptic olfactory or other sensory inputs in these tissues (Lu et al. 2009). In previous studies, ORs showing high expression in non-antennal tissues were expressed in antennae as well (Leal et al. 2013; van Schooten et al. 2020). An et al. (2016) identified highly expressed OR genes in *A. lucorum* non-antennal tissues, in which AlucOR20 showed high expression in wings and AlucOR27 in the abdomen, while expression of these two ORs in antennae was not detected. Similarly, Koutroumpa (2021) showed that 5 SlitORs were expressed only in the leg/palp of *S. littoralis*. We observed that OR30C expression from the head and OR60C from the wings was highly tissue-specific and this was not expressed in the antennae. OR60C is within the 9-exon subfamily that generally detects cuticular hydrocarbons (Pask et al. 2017), and we speculate that it performs a similar function in *C. fusciceps*.

The expression of ORs in non-antennal tissues implies that fig wasps may expand their chemosensory system for precise identification of their specific host fig species. Though ORs in the antennae act primarily in olfaction, the ORs in non-antennal tissues may have an additional role in chemical sensing. Alternatively, since the *Drosophila* wing was known to be a taste organ for many years (He et al. 2019) and during the evolution of flying insects from terrestrial organisms, the OR family evolved from the gustatory receptor (GR) family (Robertson et al. 2003; Wicher and Miazzi 2021), the remnants of GR sensing may be the manifestation of OR expression in non-antennal tissues. Later there may have been a great expansion of OR genes (Missbach et al. 2014; Brand et al. 2018; Thoma et al. 2019). He et al. (2019) reported that in *Drosophila* a candidate ionotropic pheromone receptor (IR52a) is expressed in the chemosensory sensilla of the wing. They also found that the sensilla on the wing margin express many genes including ionotropic receptors (IRs), GRs, and olfactory binding proteins (OBPs) associated with pheromone and general odor perception. Non-antennal IRs were found in the labella, pharynx, legs, and wings of *Drosophila* (Joseph and Carlson 2015; Sánchez-Alcañiz et al. 2018). Considering the extremely short life span of the fig wasp (1–2 days), the extra-antennal expression of OR receptors might help the tiny fig wasp to detect host-specific odors quickly while flying in highly variable odorscapes.

Antennal morphology is a result of various selection pressure (Elgar et al. 2018), where the size and structure of the antennae often correlates with increased sensitivity to olfactory stimuli. Antennal morphology may also reflect constraints imposed by the physical environment (Hansson and Stensmyr 2011). In the case of the fig pollinator, an increase in the size of the antennae would affect the entry of the wasp into the fig microcosm through the tiny ostiole; antennae break off and get damaged during this entry process. Hence, to compensate for the tiny size of the fig pollinator antennae, non-antennal tissues may extend their role in the precise sensing of the scents. Consequently, non-antennal tissues such as the abdomen or ovipositor may guide the wasps towards appropriate egg-laying or pollination sites within the dark fig microcosm interior even in the absence of antennae. Also, due to the tiny size of the wasp, non-antennal tissue expression may expand the surface area of tissues bearing ORs to increase olfactory sensitivity both at long and short distances.

Upon comparison of south and northeast Indian wasps regarding OR expression patterns, we observed a significant variation between these regions, not only in antennal ORs but also in all non-antennal ORs. In parallel with the OR variation, we observed variation in the volatile profile of the corresponding *F. racemosa* samples between south and northeast India during the pollen receptive phase (Additional file 1: Table S5) (Nongbri and Borges, unpublished data). A total of 65 compounds under three main groups of volatile chemicals were identified out of which 34 were found to be in common between the two regions (Nongbri and Borges, unpublished data). Around 15 volatiles were specific to south India and 16 were specific to the northeast region. This region-specific variation in the volatile profile (Additional file 1: Table S4) could be the reason for region-specific variation in OR gene expression. This question can be definitively answered only by deorphanizing the ORs.

Insects show high variation in their chemosensory genes to support rapid adaptation of odorant detection capacities (Andersson et al. 2015). Plants may also modify their VOC profiles according to the biotic and abiotic changes of the habitat (Holopainen and Gershenzon 2010; Ninkovic et al. 2020; Picazo-Aragonés et al. 2020). These joint forces may explain region-specific variation in VOC profiles and OR expression within plant–insect interactions Advances in multi-omics approaches provide a better understanding of how gene expression is regulated in response to different environmental conditions, both over short-term and long evolutionary timescales. Our results illustrate that OR evolution may lead to prominent differences in olfactory sensing at the population level within species. Persistent changes in gene expression occurring at short evolutionary scales can support cellular adaptation to environmental changes and might also trigger longer-term adaptations (López-Maury et al. 2008). These kinds of studies will also help in important future research focused on unraveling the role of insect olfactory plasticity in response to changing odorscapes since insects also show good plasticity in terms of adjusting to the rapid changes in the environment through learning (Conchou et al. 2019). Investigations with non-model insects will give insight into unexplored modes of plasticity in their olfactory receptor systems and physiological responses. Our study has also provided evidence of non-antennal expression of ORs and adds to the burgeoning data on such ectopic expressions whose function is not yet deciphered.

## Materials and Methods

### Fig pollinator collection and dissection

*Ficus racemosa* trees located in and around the Indian Institute of Science campus, Bangalore, south India, and Shillong, northeast India, were used to collect pollinator wasps. The fig bunches were enclosed with nylon mesh bags in their pre-pollination phase to prevent unregulated oviposition by fig wasps. Pollinating fig wasps, *Ceratosolen fusciceps* were introduced singly into figs during the pollen receptive stage of the syconium. The figs were allowed to mature, and the female pollinator offspring from a single foundress female were collected and immediately placed in RNA-later solution and stored at –20°C. The fig wasps stored in RNA-later were dissected to separate their tissues such as the head, thorax, abdomen, antennae, ovipositor, legs, and wings. The dissections were carried out under the microscope for the precise separation of tissues in the presence of RNA-later. A total of 14 samples of 7 tissues each for south and northeast India were taken for RNA extraction.

### Total RNA extraction

RNA-later was removed from the dissected tissues and the tissues were quickly washed with 1X PBS. The washed tissues were frozen under liquid nitrogen, mixed with cold TRIzol (Life Technologies), and homogenized using a TissueLyser II (Qiagen). The lysate was then purified using the DirectZol kit (Zymoresearch) following the manufacturer’s protocol. Additional on-column DNAseI treatment was given to remove any traces of genomic DNA according to the manufacturer’s guidelines. Total RNA bound to the membrane was eluted in RNase-free water. The quality and quantity of isolated RNA were analyzed using a nanophotometer (IMPLEN). The integrity of total RNA was assessed in a bio-analyzer (Agilent 2000, Agilent Technologies, USA) using an RNA 6000 Nano Lab Chip (Agilent Technologies, USA). The RNA integrity (RIN) was calculated by considering 18s and 28s ribosomal RNA ratios and baseline correction factors. The samples with >7 RIN values were considered for further analysis.

### cDNA library prep

Good quality RNA at the required concentration was used to synthesize a cDNA library using NEB Next Ultra Directional RNA library prep. RNA (1 μg) was taken for mRNA isolation, fragmentation, and priming. The fragmented and primed mRNA was further subjected to first-strand synthesis in the presence of Actinomycin D (Gibco, Life Technologies, CA, USA) followed by second-strand synthesis. The double-stranded cDNA was purified using HighPrep magnetic beads (Magbio Genomics Inc, USA). The purified cDNA was end-repaired, adenylated, and ligated to Illumina multiplex barcode adapters according to the NEB Next Ultra Directional RNA Library Prep Kit protocol. The adapters used in the study were the Illumina Universal Adapter:

5’AATGATACGGCGACCACCGAGATCTACACTCTTTCCCTACACGACGCTCTTCC GATCT-3’ and Index Adapter:

5’-GATCGGAAGAGCACACGTCTGAACTCCAGTCAC[INDEX] ATCTCGTATGCCGTCTTCTGCTTG-3’.

The adapter-ligated cDNA was purified using HighPrep beads and was subjected to 15 cycles of Indexing-PCR (37°C for 15 mins followed by denaturation at 98°C for 30 secs), and cycling (98°C for 10 sec, 65°C for 75 sec and 65°C for 5 mins) to enrich the adapter-ligated fragments. The final PCR product (sequencing library) was purified with HighPrep beads, followed by a library-quality control check. The Illumina-compatible sequencing library was initially quantified by Qubit fluorometer (Thermo Fisher Scientific, MA, USA) and its fragment size distribution was analyzed on Agilent 2200 TapeStation.

### RNA-Sequencing and preprocessing

The cDNA library fragment size ranged from 300 bp to 700 bp. As the combined adapter size is approximately 120 bp, the effective insert size is 180 bp to 580 bp. Thus, the resultant cDNA library had enough concentration and was suitable for paired-end (150*2) sequencing using an Illumina HiSeq 4000 platform to get the desired amount of sequencing data. The raw sequencing data were checked for quality using FastQC (Andrews 2010) and were pre-processed, which included removing adapter sequences and low-quality bases. Raw reads were processed by “Cut Adapt” for adapters (Martin 2011) and low-quality bases trimming towards 3’-end using Trimmomatic from Trinity. An average of 23.87 (South India) and 21.23 million (Northeast India) preprocessed reads were used for downstream analysis.

### Genome-guided transcriptome assembly

Pre-processed RNA-seq reads from seven fig wasp tissues across two regions were mapped separately to the *C. fusciceps* genome with mapping rates of 73% (south India) and 60% (northeast India) for the pooled tissues of each region. STAR (2.7.0a) (Dobin et al. 2013). These two mapping files were used to assemble genome-guided Trinity assemblies (Haas et al. 2013) for the two regions (v 2.8.5) The initial number of transcripts obtained were 262904 and 255260 for south and northeast India with contig N50 of 1323 and 1227 base pairs, respectively. These two sets of transcripts were combined and the transcripts with 100% sequence identity were removed which resulted in 334449 transcripts with contig N50 of 1420 base pairs. To avoid the removal of highly similar paralogs, as these were often found in olfactory receptors, a lower redundancy cutoff was not chosen. Instead, further filtering was done based on the alignment of these transcripts to the high-quality genome and the annotation of these transcripts. Mainly longer, protein-coding and known protein-domain containing transcripts were prioritized over other transcripts. ‘Transdecoder’ (Gotoh 2000) was used to translate the predicted transcripts and hmmscan (Mistry et al. 2013) was used to detect the presence of any known protein domains from the Pfam protein family database (Sonnhammer et al. 1997; Finn et al. 2015). All these 3,34,449 transcripts were aligned to the *C. fusciceps* draft genome (Supplemental Table S1). using SPLAN 2.3.2a (Iwata and Gotoh 2012). This aligner predicts approximate ‘gene’ regions/clusters where multiple transcripts align. Each such ‘gene’ region was examined individually, and preference was given to the longest transcripts and the remaining transcripts were compared with the longer transcript. The following criteria were used to select one or few representative transcripts per gene cluster: If one transcript was found nested within another transcript (according to the alignment boundaries taken from the alignments with the genome), the shorter one was removed if the alignment score was lower and or no protein domain was found within. In the case of partial overlap amongst transcripts with a minimum of 50% overlap to the longest transcript, the one without the protein domain was removed. In case both or neither of the partially overlapping transcripts coded for a valid protein domain, then the one with the lesser alignment score was removed. In this way, few alternatively spliced transcripts that were sufficiently different from the longest transcripts per gene were also retained. The number of finally retained transcripts was 58,076. Out of these 17,746 were protein-coding and 14,417 contained known protein domains. RSEM with Bowtie2 (Langmead and Salzberg 2012) was used for mapping reads to the transcriptome assembly and further quantification of expression. Around one thousand transcripts without any read mapping were removed. The mapping was used to create an expression matrix for all 14 tissues from 2 different regions containing FPKM/TPM values. On average South Indian reads had 62.2% mapping and Northeast Indian reads had 52.1% mapping. Approximately an average of 10% of reads were not mapped per tissue per location.

### Gene Annotation

UniProt (Bateman et al. 2020) and KAAS (Moriya et al. 2007) were used for functional annotation of the transcripts. Clustered transcripts were annotated using the homology approach to assign functional annotation using BLAST (Camacho et al. 2009) against “Insecta” data from the Uniprot database containing 2,883,368 protein sequences. Transcripts were assigned with a homolog protein from other organisms if the match was found at an e-value less than e-5 and a minimum identity of greater than 30%.

### OR annotation

The putative OR transcripts with the 7tm_6 domain were identified as potential Olfactory Receptors with the help of hmmscan against the Pfam database. This resulted in 74 putative OR transcripts. To corroborate the results, the InsectOR web server was used on the combined transcriptome containing 3,34,449 unique transcripts. InsectOR performs OR gene prediction directly from the genome/transcriptome without prior prediction of proteins from another tool and is more sensitive than a few other genome annotation tools (Karpe et al. 2021). This resulted in the identification of 514 protein sequences with the 7tm_6 domain. These were further filtered based on the presence of the identified transcripts in the final non-redundant transcript assembly containing the 58K chosen transcripts. This resulted in 74 ORs as before. These 74 transcripts were manually studied, 3 transcripts were found to produce identical amino acid sequences to another sequence and a few others were found to be arising from the same genomic location with minor differences. Finally, 63 manually curated high-quality OR transcripts were used for further studies (Additional file 2).

### Phylogenetic analysis

Well-curated OR protein sequences from *C. fusciceps* (this study), *C. solmsi* (Zhou et al. 2015), *M. demolitor* (Zhou et al. 2015), *N. vitripennis* (Robertson et al. 2010)*, A. mellifera* (Robertson and Wanner 2006)*, D. novaeanglie* (Robertson and Wanner 2006) and *H. saltator* (McKenzie et al. 2014) were collected and partial sequences (<200 amino acids) were removed. Chosen sequences were aligned using MAFFT (v7.123b, E-INS-i strategy, JTT200 matrix, 1000 iterations). The alignment was trimmed using trimAl (‘automated1’ option). A maximum likelihood (ML) based phylogenetic tree was reconstructed using RAxML (v7.4.2, PROTCATJTTF matrix, 100 rapid bootstraps, seven olfactory receptor-coreceptor (Orco) sequences as outgroup). This tree was used as a guide for the second iteration of the alignment by MAFFT (Katoh et al. 2002; Katoh and Standley 2013). The second alignment was trimmed using the trimAl option ‘gappyout’ (Capella-Gutierrez et al. 2009). The refined phylogenetic tree was reconstructed again with RAxML (Stamatakis 2006) with similar parameters. (iTOL) v3 (Letunic and Bork 2007, 2016) was used for tree visualization. The tree was annotated and divided into subfamilies /clades with the help of an existing hymenopteran OR tree.

### OR expression matrix generation

To find the expression profile of 63 ORs, RNA-seq reads were mapped to the transcriptome using Bowtie2 (Langmead and Salzberg 2012) and RSEM (Li and Dewey 2011). An average of 62.2% and 52.1% mapping rates were obtained for two regions because the transcriptome was assembled with 60–70% of the total reads that mapped to the genome. Also, an average of 10% of reads was not mapped per tissue per region. The mapped transcripts were used to create an OR expression matrix for all 14 tissues from the 2 regions containing TPM (transcripts per million) values (Additional file 3) and plotted in the form of a heatmap for easy comparison.

### qPCR Analysis

RNA was isolated as explained previously. According to the manufacturer’s protocol, the total RNA was converted into cDNA using Prime Script RT Reagents (Takara). In brief, approximately 500 ng of RNA from each sample was taken for cDNA synthesis and the first strand of cDNA was synthesized using universal oligo dT primers. The synthesized cDNA was stored at -20°C. The primers were designed (Additional file 1: Table S6) for selected genes using Primer 3 Plus online primer design software considering the exon and coding regions of the transcripts. The designed primers were analyzed for their specificity by In-Silico PCR in UCSC *In-silico* PCR online bioinformatics tool and the primer characteristics were analyzed in a multiple primer analyzer (Thermo Scientific, USA) for the possibility of primer dimer formation. The primer sequences that passed all the quality criteria were processed for synthesis on a 10 nm scale and purified by HPLC. The final set of primer sequences is listed in (Additional file 1: Table S6). The synthesized primers were validated for their specificity using pooled cDNA from all the tissues of the south and northeast regions. In brief, the 1μ sample were pooled and diluted to the final concentration of 10 ng^-μ^ and 1 L was used for each μ qPCR reaction with picomols (pM) of primer concentration. The primers which showed a good amplification curve with a single melt curve and desired specific product size on agarose gel were used for further relative quantification by qPCR. The expression levels of selected genes were analyzed using SYBR Green chemistry (Brilliant II SYBR Green qPCR master mix (Agilent Technologies, USA) in the Stratagene mx3005P instrument (Agilent Technologies, USA). The amplification cycling conditions were as follows: initial denaturation for 95°C for 10min followed by 40 cycles of 95°C for 30 sec, 60°C for 30 sec. The dissociation curve analysis was performed after amplification for primer specificity; the conditions were as follows: 95°C for 1min, 55°C, for 30 sec, and 0.2°C/sec increment up to 95°C (continuous fluorescence collection from 55–95°C). The mean Ct value of technical replicates was used to calculate the relative expression level of genes. The relative quantification of genes was analyzed using standard 2^-ΔΔ^ as described by Pfaffl (2001). Fold-change values in log base 2 values from 5 biological replicates of each tissue were compared across sites by t-tests. P values are provided in Additional file 1: Table S7. Beta-actin was used as a reference gene to normalize the qPCR experiment after comparing it with RPS18. The subsets of OR genes for qPCR analysis were selected in such a way that each selected OR showed upregulated expression in one region (south or northeast India) and downregulated expression in another region (south or northeast India). In addition, ORs were selected that showed equal levels of expression. Accordingly, the following ORs were selected (showing opposite patterns of expression at the two sites): OR32 and the OR52 from the abdomen, OR8 and OR62 from the antenna, OR3C, OR40F, and OR64C from the ovipositor, OR46 and OR60C from wings, OR7C from the head, OR32 and OR40F from legs, OR32 and OR52 from the thorax, and the following showing nearly equal expression at the two sites: OR64C from the abdomen, OR17F from antennae and wings, OR66 from the head.

## Declaration

### Ethics approval and consent to participate

Ethics approvals are not applicable to this study.

### Data Availability

The data generated and analyzed in this study are included within the manuscript and supplementary data. All raw sequencing data generated in this study have been submitted to the NCBI SRA (https://submit.ncbi.nlm.nih.gov/subs/sra/SUB11600929). The raw RNA-seq reads data generated in this study have been submitted to the NCBI Bio Project (PRJNA853513) database under accession number SUB11600929.

## Acknowledgments

This work was funded by the Department of Biotechnology (DBT), under the project entitled, ‘Chemical Ecology of the Northeast Region (NER) of India: A collaborative program linking NER and Bangalore Researchers’ (DBT-NER/Agri/24/2013 dated 30/03/2013). We also acknowledge support from the DBT-IISc partnership program, and DST-FIST. Sushma Krishnan is grateful for an EMBO Short-term fellowship that enabled the transcriptome work done at Max-Plank Institute of Chemical Ecology, Jena. Snehal Karpe is grateful for CSIR Shyama Prasad Mukherjee Fellowship and NCBS Bridging Postdoctoral Fellowship. We thank G. Yathiraj for the fig collection from south India and Anusha Kumble for her great help in wasp sample collection and dissection.

## Authors contribution

Conception, R.M.B., and S.K.; Sample collection, S.K. and L.B.N.; Dissection and RNA isolation, S.K.; Transcriptomics, S.K. H.K and E.G.; Genome assembly, H.K.; Genome guided transcriptome assembly, S.D.K.; OR annotation, Phylogeny, Expression analysis, S.K., S.D.K. and E.G.; qPCR analysis, S.K.; Writing, S.K., S.D.K., E.G., B.S.H., and R.M.B.; Supervision, S.R., B.S.H., E.G. and R.M.B.. The authors read and approved the final manuscript.

## Supplementary Information

Supplementary information is available in the additional file.

### Additional File 1

**Figure. S1.**
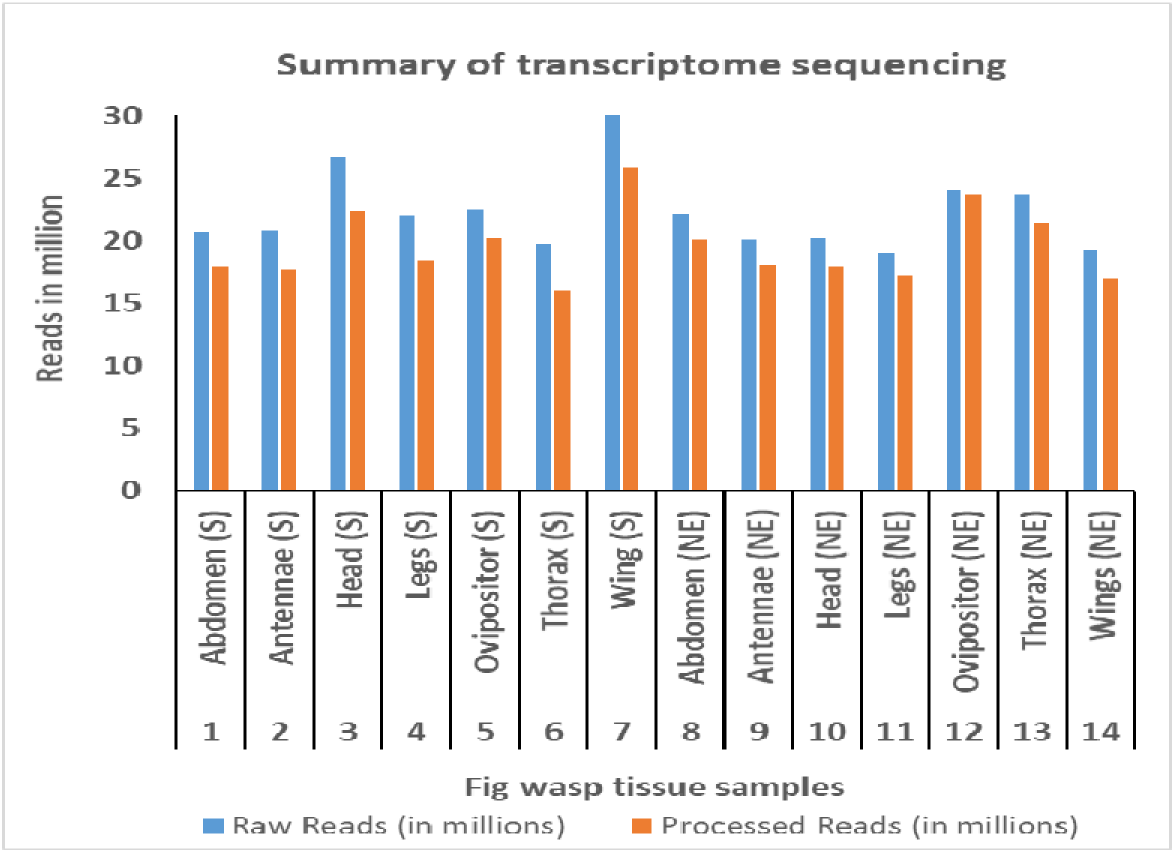
Bar chart indicating raw and processed reads obtained from fig wasp transcriptome sequencing. Illumina Hi-Seq, 150 X 2 paired-end sequencing was used to sequence 7 tissues each from South India (S) and Northeast India (NE).

**Figure S2.**
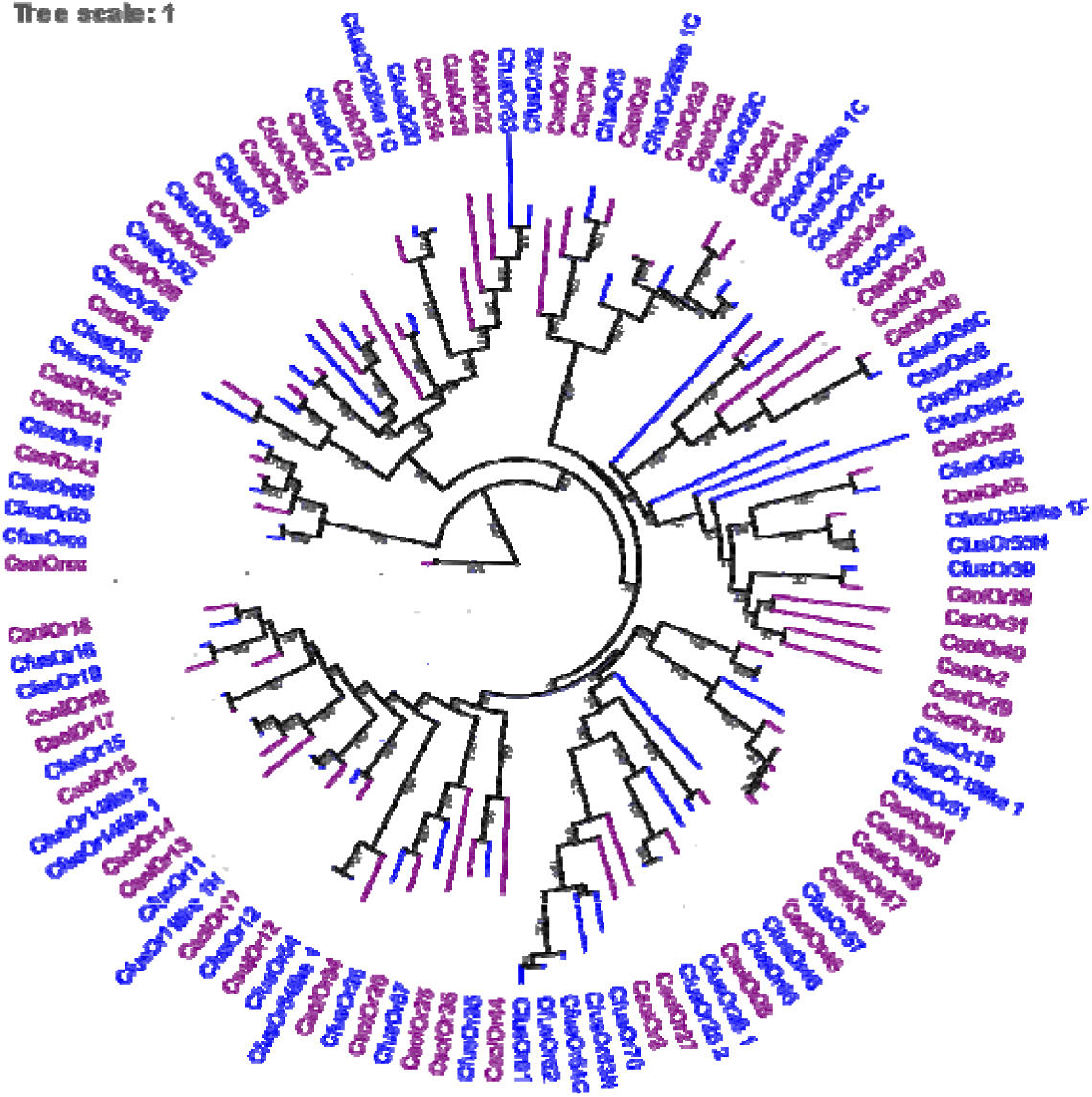
Phylogenetic tree reconstruction with *C. fuscicpes* and *C. solmsi* olfactory receptors The branches are color coded as per species - *C. fusciceps* - Blue, *C. solmsi* – Purple.

**Table S1.** Genome Hybrid assembly statistics

| Scaffolds |  | Contigs |  |
| --- | --- | --- | --- |
| Scaffolds Generated: | 1286 | Contigs Generated: | 1497 |
| Maximum Scaffold Length (bp): | 1,11,87,473 | Maximum Contig Length (bp): | 59,78,652 |
| Minimum Scaffold Length (bp): | 1001 | Minimum Contig Length (bp): | 165 |
| Average Scaffold Length (bp): | 185043 | Average Contig Length (bp): | 158841 |
| Median Scaffold Length (bp): | 4129 | Median Contig Length (bp): | 1013 |
| **Total Scaffolds Length (bp):** | **23,79,66,267** | **Total Contigs Length (bp):** | **23,77,85,791** |
| Scaffolds >= 100 bp: | 1286 | Contigs >= 100 bp: | 1497 |
| Scaffolds >= 200 bp: | 1286 | Contigs >= 200 bp: | 1496 |
| Scaffolds >= 500 bp: | 1286 | Contigs >= 500 bp: | 1492 |
| Scaffolds >= 1 Kbp: | 1286 | Contigs >= 1 Kbp: | 1490 |
| Scaffolds >= 10 Kbp: | 232 | Contigs >= 10 Kbp: | 346 |
| Scaffolds >= 1 Mbp: | 57 | Contigs >= 1 Mbp: | 72 |
| **N50 value:** | **41,21,315** | **N50 value:** | **22,03,451** |

**Table S2.** BUSCO analysis of transcriptome assembly

|  | **Hymenoptera** | **Insecta** |
| --- | --- | --- |
| Complete BUSCOs (C) | 3002 | 1358 |
| Complete and single-copy BUSCOs (S) | 2388 | 1068 |
| Complete and duplicated BUSCOs (D) | 614 | 290 |
| Fragmented BUSCOs (F) | 450 | 77 |
| Missing BUSCOs (M) | 963 | 223 |
| Total BUSCO groups searched (N) | 4415 | 1658 |
| Result Summary | C:68.0% [S:54.1%,<br>D:13.9%], F:10.2%,<br>M:21.8%, N:4415 | C:81.9% [S:64.4%,<br>D:17.5%], F:4.6%,<br>M:13.5%, n:1658 |

**Table S3.**
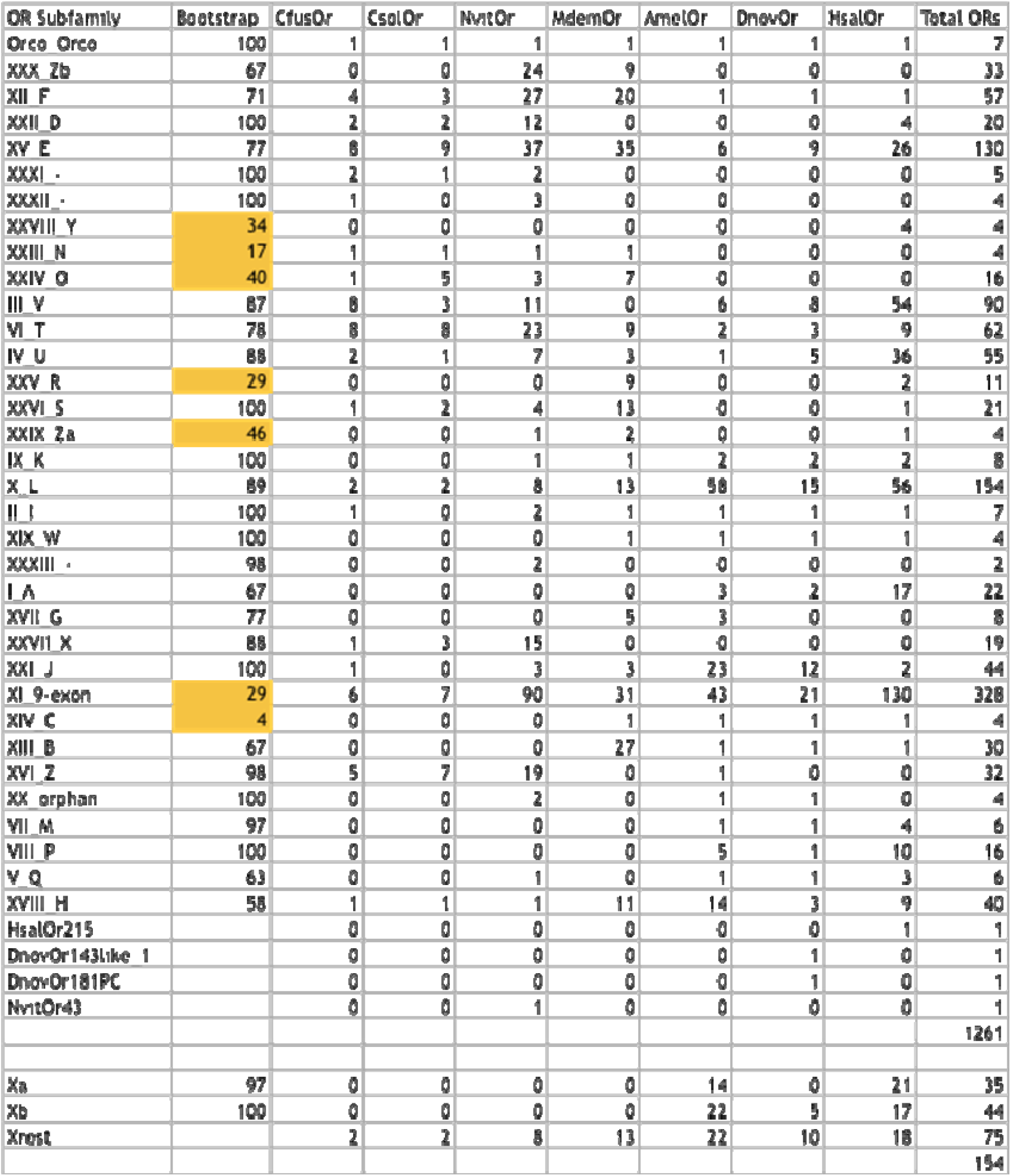
Detailed information about the hymenopteran OR phylogenetic tree

**Table S4.**
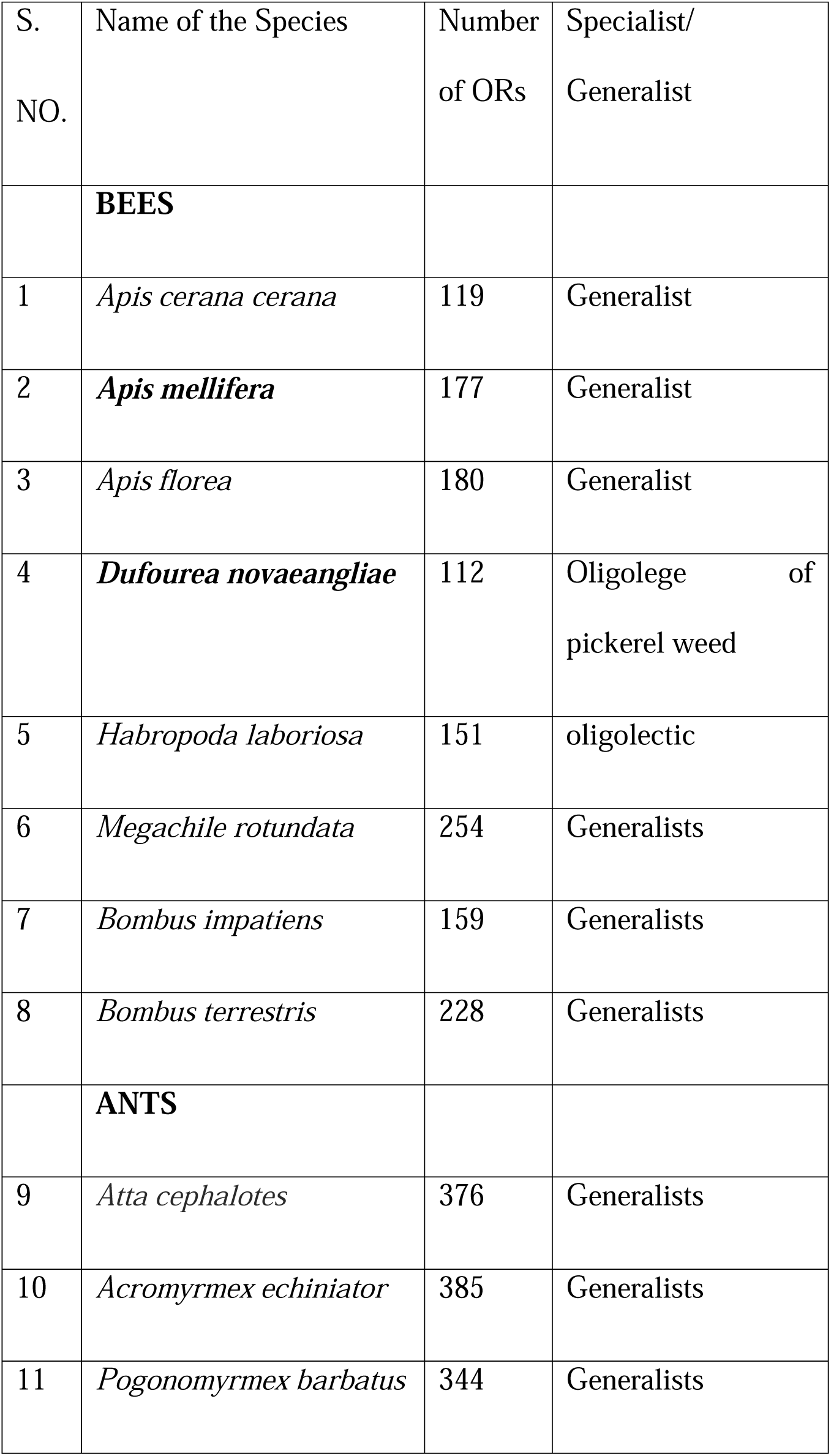
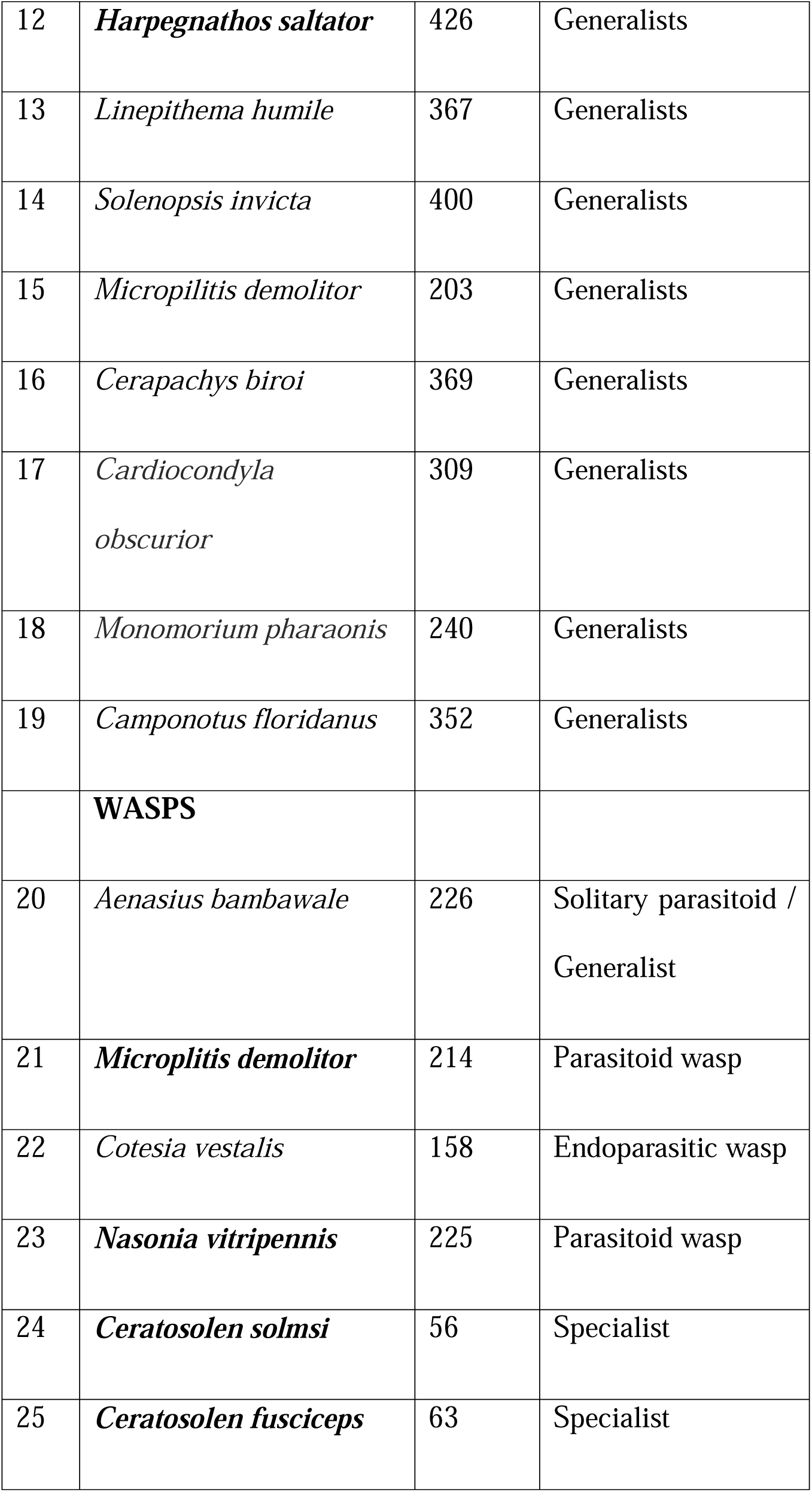
Number of annotated ORs in specialist and generalist Hymenoptera

| S.<br>NO. | Name of the Species | Number<br>of ORs | Specialist/<br>Generalist |
| --- | --- | --- | --- |
|  | **BEES** |  |  |
| 1 | *Apis cerana cerana* | 119 | Generalist |
| 2 | *Apis mellifera* | 177 | Generalist |
| 3 | *Apis florea* | 180 | Generalist |
| 4 | *Dufourea novaeangliae* | 112 | Oligolege of<br>pickerel weed |
| 5 | *Habropoda laboriosa* | 151 | oligolectic |
| 6 | *Megachile rotundata* | 254 | Generalists |
| 7 | *Bombus impatiens* | 159 | Generalists |
| 8 | *Bombus terrestris* | 228 | Generalists |
|  | **ANTS** |  |  |
| 9 | *Atta cephalotes* | 376 | Generalists |
| 10 | *Acromyrmex echiniator* | 385 | Generalists |
| 11 | *Pogonomyrmex barbatus* | 344 | Generalists |
| 12 | *Harpegnathos saltator* | 426 | Generalists |
| 13 | *Linepithema humile* | 367 | Generalists |
| 14 | *Solenopsis invicta* | 400 | Generalists |
| 15 | *Micropilitis demolitor* | 203 | Generalists |
| 16 | *Cerapachys biroi* | 369 | Generalists |
| 17 | *Cardiocondyla*<br>*obscurior* | 309 | Generalists |
| 18 | *Monomorium pharaonis* | 240 | Generalists |
| 19 | *Camponotus floridanus* | 352 | Generalists |
|  | **WASPS** |  |  |
| 20 | *Aenasius bambawale* | 226 | Solitary parasitoid /<br>Generalist |
| 21 | *Microplitis demolitor* | 214 | Parasitoid wasp |
| 22 | *Cotesia vestalis* | 158 | Endoparasitic wasp |
| 23 | *Nasonia vitripennis* | 225 | Parasitoid wasp |
| 24 | *Ceratosolen solmsi* | 56 | Specialist |
| 25 | *Ceratosolen fusciceps* | 63 | Specialist |

**Table S5.** Proportional abundance (%) of VOC’s from receptive (B-phase) of *Ficus racemosa* trees (Means ± SD, n=19 for SI, n=17 for NE; RI=Retention Index) from North-east (Meghalaya) and South India (Bangalore)

| S.NO | Compounds | R I | South India (SI) | Northeast India (NE) |
| --- | --- | --- | --- | --- |
| **I** | **Fatty acid derivatives** |  |  |  |
| 1 | (Z)-3-hexen-1-ol | 856 | 2.53±1.41 | 0.05±0.20 |
| 2 | Hexanol | 868 | 0.19±0.20 | - |
| 3 | 1-octen-3-ol | 978 | - | 0.33±0.90 |
| 4 | 3-octanone | 979 | - | 0.22±0.92 |
| 5 | octan-3-ol | 997 | - | 0.02±0.10 |
| 6 | hexyl acetate | 1007 | 0.59±0.97 | 0.23±0.85 |
| 7 | (Z)-3-hexenyl acetate | 1009 | 54.41±22.54 | - |
| 8 | nonanal | 1105 | 6.43±8.56 | 1.79±3.55 |
| 9 | decanal | 1204 | 0.28±0.54 | 0.19±0.76 |
| 10 | (Z)-hex-2-en-1-yl benzoate | 1573 | 0.11±0.21 | - |
|  | **Total** |  | **64.53±34.44** | **2.83±7.27** |
| **II** | **Aromatics (Shikimic acid pathway)** |  |  |  |
| 11 | Benzaldehyde | 953 | 0.57±1.66 | 1.96±2.35 |
| 12 | benzyl alcohol | 1034 | 2.47±4.33 | 3.35±4.74 |
| 13 | 2-phenylethanol | 1111 | 0.08±0.24 | - |
| 14 | benzyl acetate | 1163 | 1.58±3.22 | - |
| 15 | methyl salicylate | 1195 | 0.20±0.87 | - |
| 16 | Indole | 1289 | 0.16±0.47 | 1.28±2.60 |
|  | **Total** |  | **5.07±10.79** | **6.59±9.69** |
| **III** | **Monoterpenes** |  |  |  |
| 17 | a-thujene | 920 | - | 0.27±0.77 |
| 18 | α-pinene | 932 | 0.21±0.41 | 0.39±0.95 |
| 19 | sabinene | 971 | - | 0.50±0.78 |
| 20 | β -pinene | 974 | - | 0.08±0.20 |
| 21 | 6-methyl-5-hepten-2-one | 988 | 1.22±1.78 | 2.09±6.26 |
| 22 | myrcene | 988 | - | 0.58±1.12 |
| 23 | 6-methyl-5-hepten-2-ol | 993 | 0.81±1.24 | 0.38±0.63 |
| 24 | Limonene | 1029 | 1.15±2.62 | 0.65±1.23 |
| 25 | 1,8-cineole | 1031 | 0.03±0.12 | 2.97±3.90 |
| 26 | (Z)-β-ocimene | 1040 | 0.50±0.61 | 0.80±1.05 |
| 27 | (E)-β-ocimene | 1049 | 20.52±20.98 | 14.98±20.51 |
| 28 | NI 1 | 1056 | - | 0.31±1.27 |
| 29 | Linalool | 1099 | 1.09±2.25 | 0.40±1.35 |
| 30 | α-terpineol | 1187 | 0.04±0.17 | - |
|  | **Total** |  | **25.57±30.19** | **24.63±40.85** |
| **III** | **Sesquiterpenes** |  |  |  |
| 31 | (E)-4,8-dimethyl-1,3,7-nonatriene | 1117 | 1.81±4.11 | 15.16±21.45 |
| 32 | Geraniol | 1256 | - | 0.22±0.83 |
| 33 | α-ylangene | 1375 | 0.02±0.05 | - |
| 34 | α-copaene | 1378 | 0.09±0.38 | 0.63±0.71 |
| 35 | Daucene | 1385 | 0.11±0.21 | 1.29±1.76 |
| 36 | NI 2 | 1383 | - | 0.91±1.44 |
| 37 | α-duprezianene | 1387 | 0.09±0.17 | 0.87±1.12 |
| 38 | 7-epi-a-cedrene | 1407 | 0.02±0.05 | 1.08±0.89 |
| 39 | cis-α-bergamotene | 1408 | 0.03±0.09 | 0.31±0.40 |
| 40 | β-funebrene | 1418 | 0.16±0.32 | 2.61±1.89 |
| 41 | β-ylangene | 1420 | 0.34±0.69 | 9.32±6.13 |
| 42 | β-duprezianene | 1421 | - | 0.57±1.37 |
| 43 | β-cedrene | 1425 | 0.01±0.05 | - |
| 44 | β-copaene | 1427 | 0.05±0.11 | 1.97±1.43 |
| 45 | β-gurjunene | 1436 | 0.02±0.08 | 1.36±2.22 |
| 46 | NI 3 | 1438 | 0.07±0.17 | - |
| 47 | α-guaiene | 1440 | - | 0.33±0.51 |
| 48 | (Z)-β-farnesene | 1450 | 0.12±0.49 | - |
| 49 | epi-prezizaene | 1454 | 0.08±0.17 | 2.25±2.11 |
| 50 | epi-zizaene | 1456 | 0.21±0.50 | 8.15±7.06 |
| 51 | (E)-β-farnesene | 1458 | 0.14±0.49 | - |
| 52 | α-acoradiene | 1460 | 0.09±0.26 | 0.79±2.42 |
| 53 | zizaene (= khusimene) | 1466 | 0.93±2.05 | 13.95±12.63 |
| 54 | NI 4 | 1468 | - | 0.95±1.81 |
| 55 | β-acoradiene | 1467 | 0.03±0.13 | - |
| 56 | 10-epi-β-acoradiene | 1480 | 0.07±0.16 | 0.52±1.05 |
| 57 | γ-curcumene | 1483 | 0.01±0.04 | 0.14±0.26 |
| 58 | ar-curcumene | 1486 | 0.00±0.02 | 0.08±0.13 |
| 59 | (E,E)-α-farnesene | 1496 | 0.03±0.14 | - |
| 60 | (Z)-γ-bisabolene | 1510 | 0.24±0.71 | - |
| 61 | α-alaskene | 1511 | - | 1.05±2.64 |
| 62 | β-curcumene | 1516 | 0.03±0.13 | 0.29±0.38 |
| 63 | β-sesquiphellandrene | 1529 | 0.03±0.10 | 0.73±0.88 |
| 64 | (E)-nerolidol | 1569 | - | 0.04±0.10 |
| 65 | zizanone | 1679 | - | 0.59±1.67 |
|  | **Total** |  | **4.83±11.87** | **65.95±74.45** |

**Table. S6.**
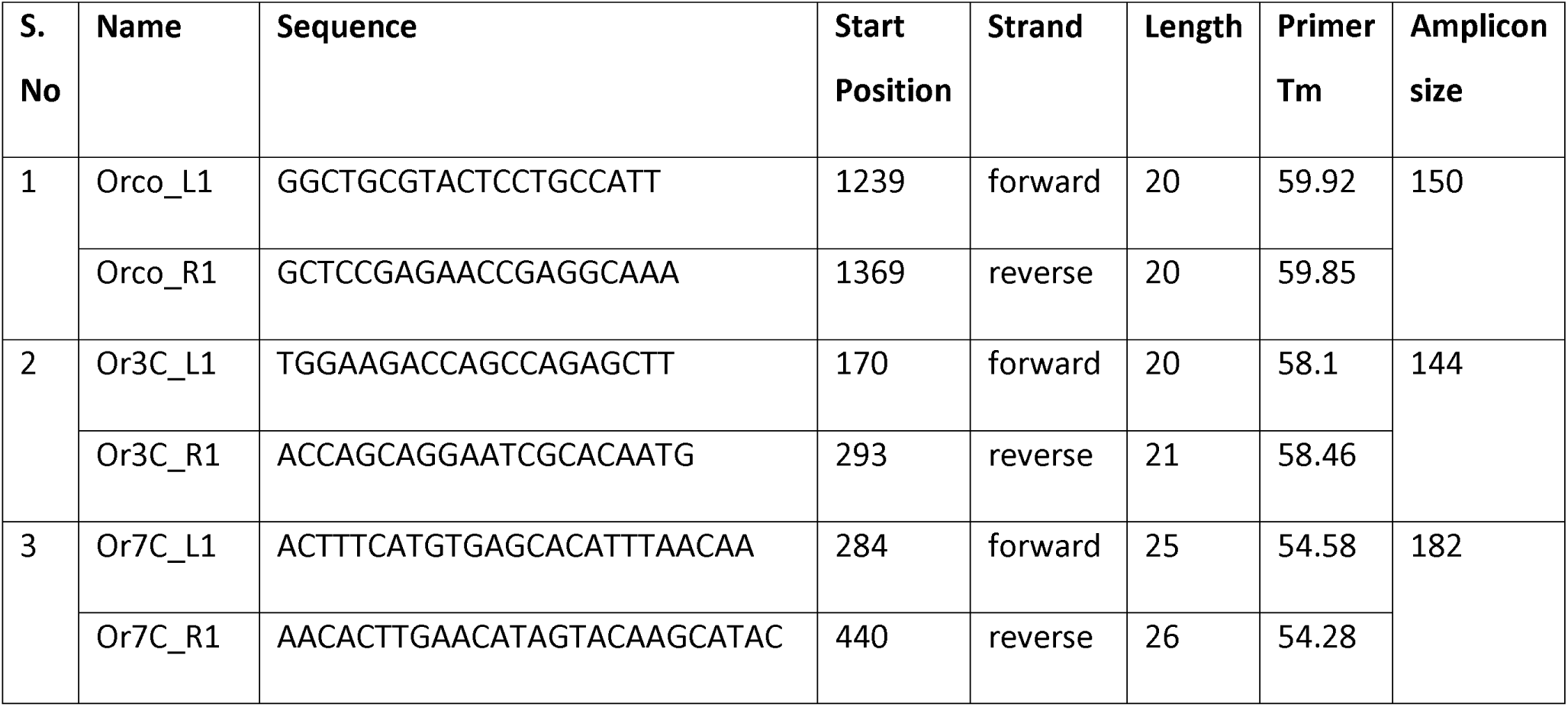
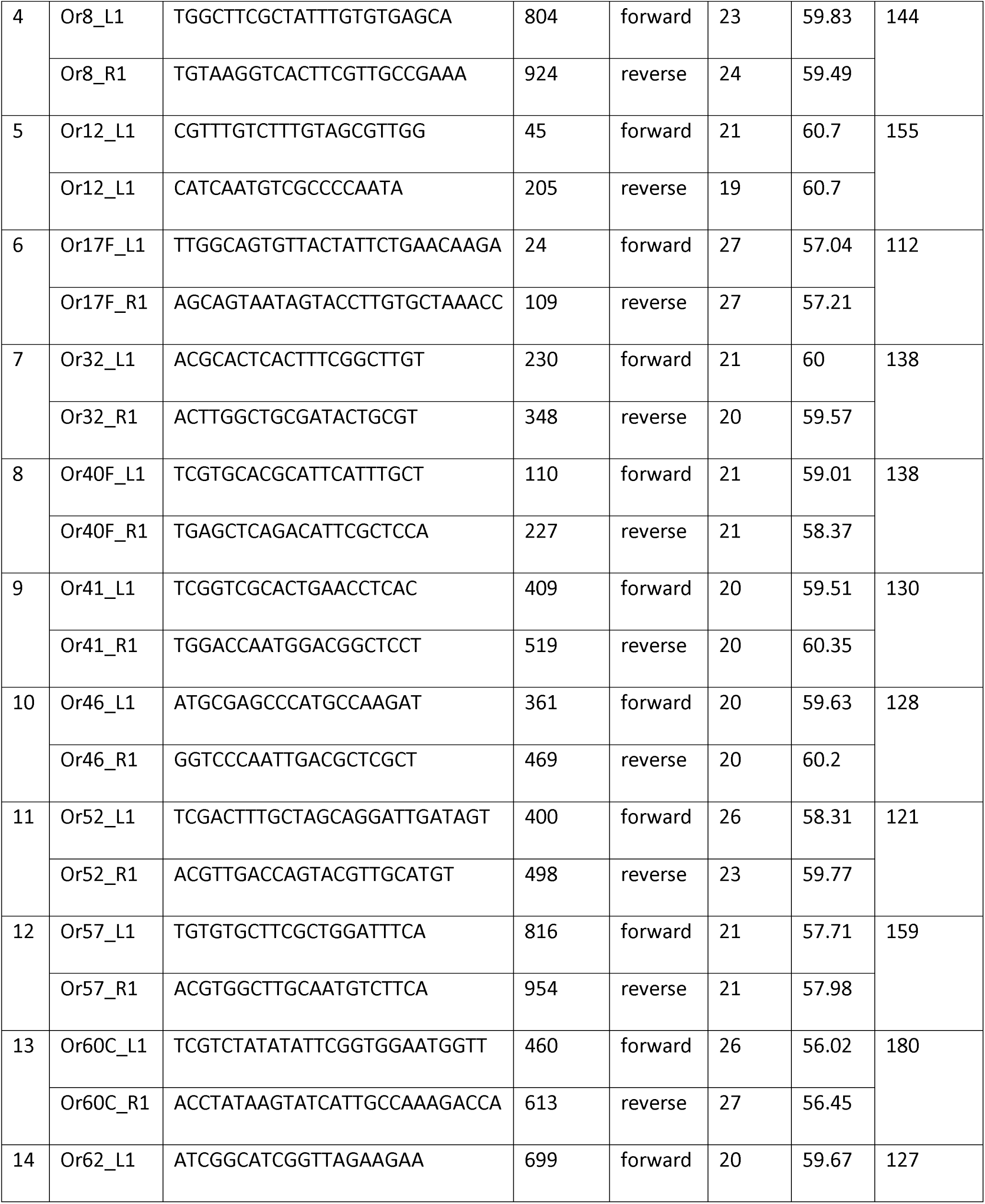
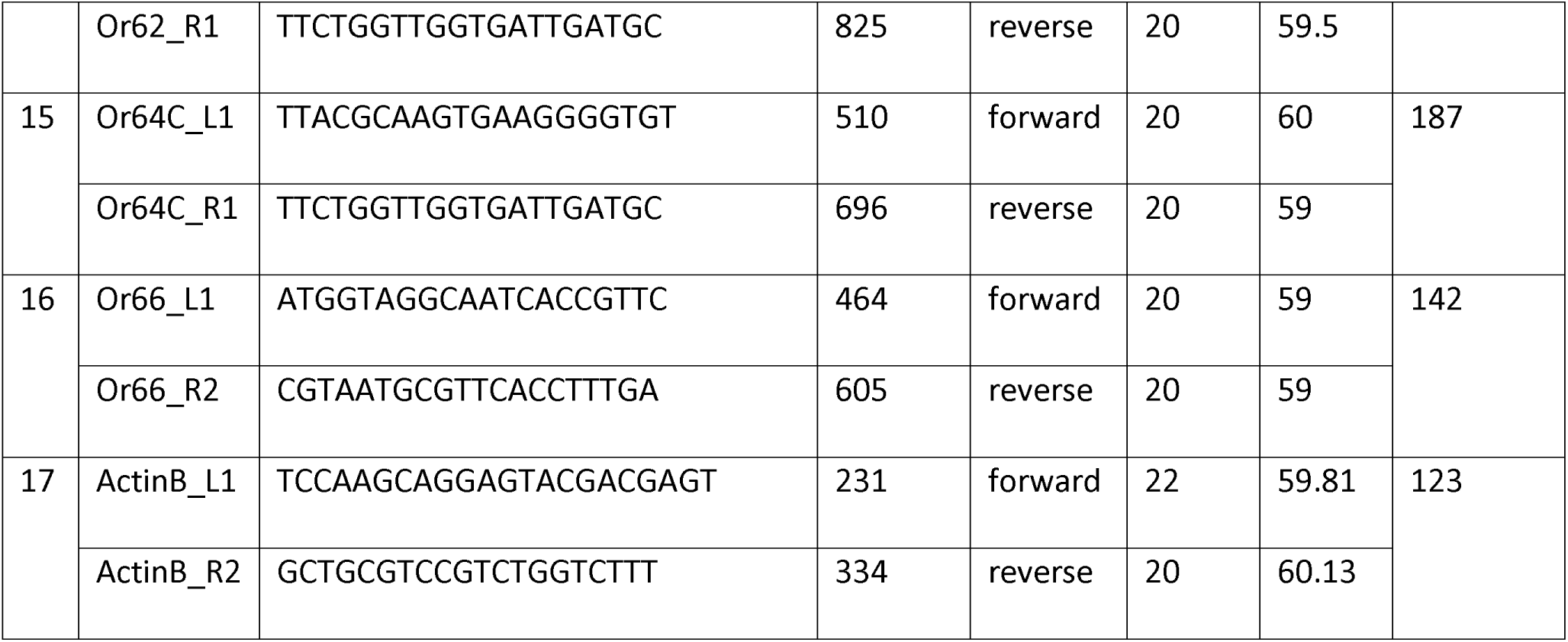
Primer sequences for OR genes

**Table S7.**
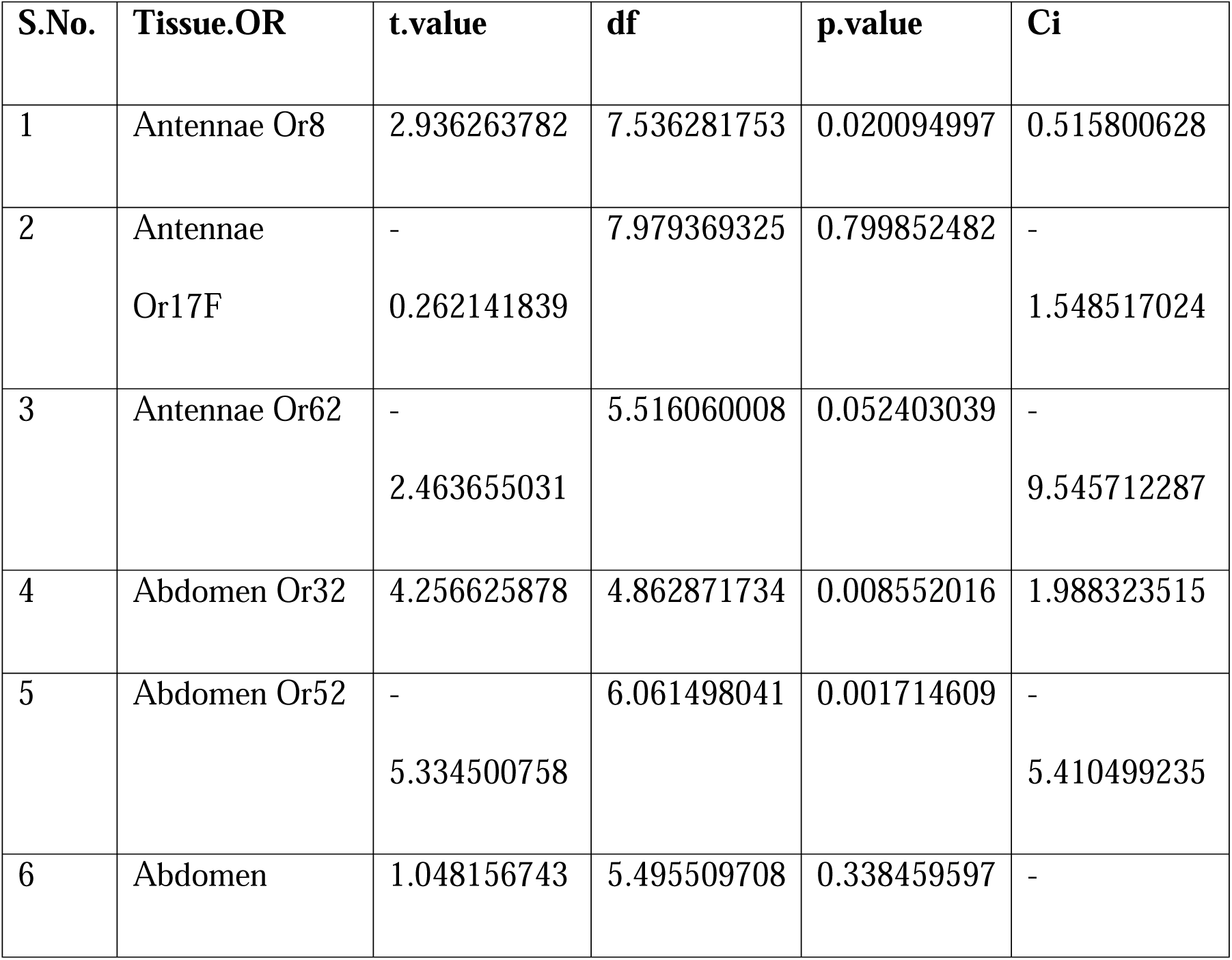
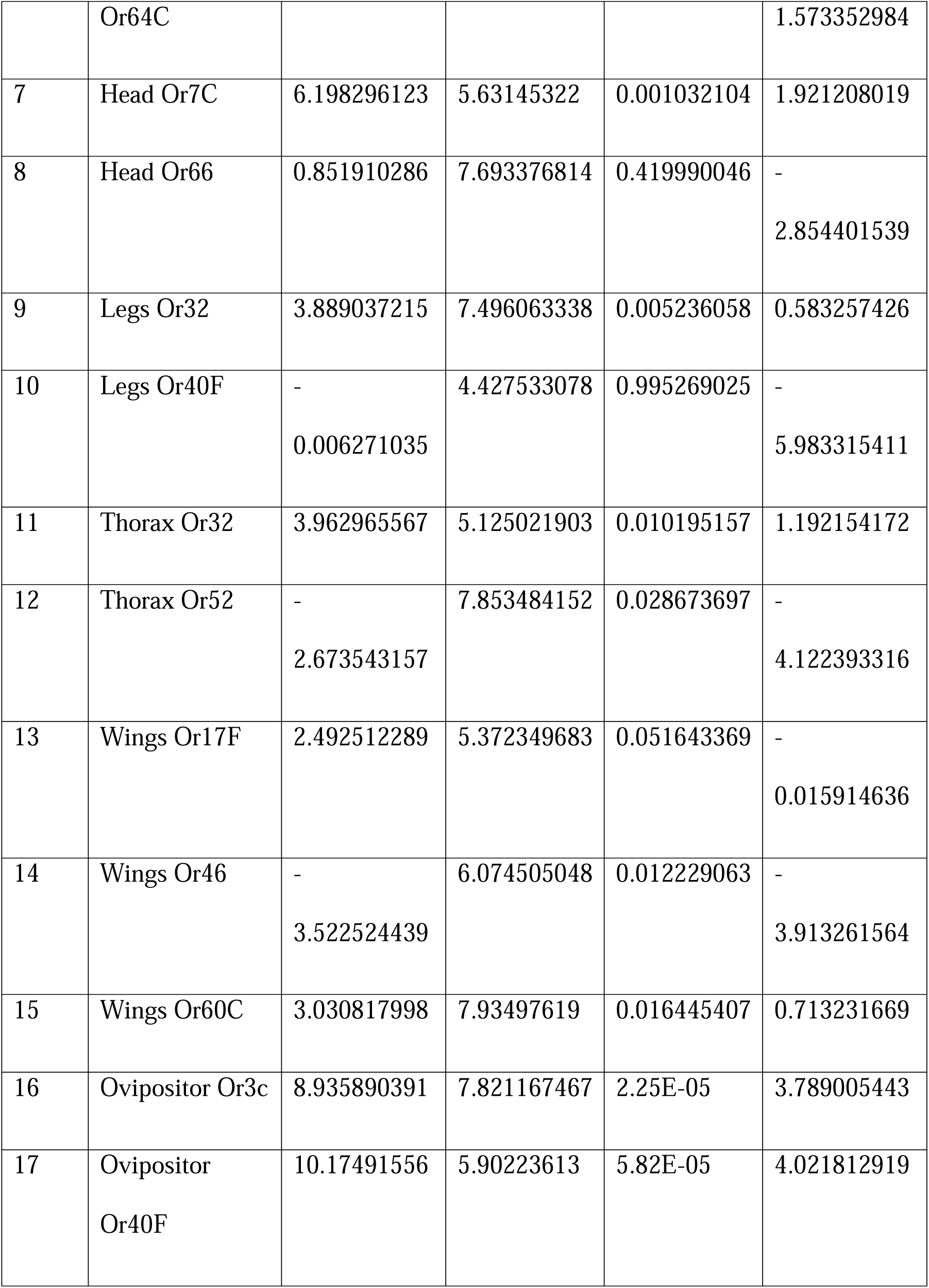
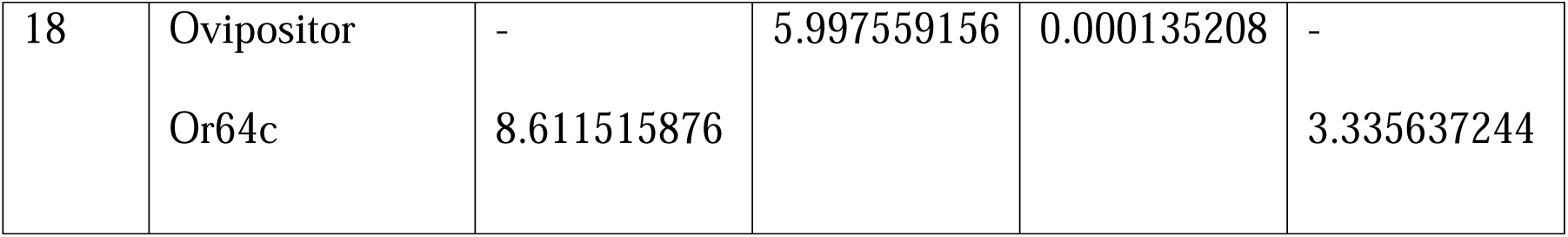
Statistical analysis of qPCR analysis of Cfus ORs

### Additional File 2

Annotation of *C.fusciceps* olfactory receptors

### Additional File 3

Expression matrix containing transcripts per million (TPM) values for *C.fusciceps* olfactory receptors

